# Novel connections between B-vitamins and microbial communities along biogeochemical gradients in a large temperate estuary

**DOI:** 10.64898/2026.02.26.707256

**Authors:** Meriel J. Bittner, Catherine C. Bannon, Elden Rowland, Gregor Luetzenburg, Erin M. Bertrand, Lasse Riemann, Ryan W. Paerl

## Abstract

As B-vitamins are organic cofactors required by prokaryotic and eukaryotic planktonic cells, their availability impacts aquatic microbial communities and associated biogeochemistry. Contrary to inorganic nutrients, measurements of B-vitamins from brackish systems are scarce and relationships between B-vitamins and plankton composition in estuaries are unclear, limiting our understanding of estuary biology in general as well as how B-vitamins are distributed and dispersed in marine systems. Here, we quantify multiple B-vitamins and their vitamers in particulate and dissolved phases, and characterize microbial community composition, across fresh to polyhaline zones of the Neuse River Estuary (NRE), North Carolina, USA. We uncover elevated concentrations of B-vitamins within the mid-estuary, Chlorophyll *a* maximum along with a unique suite of dissolved B-vitamin associated with sporadic surges in pico- and microplankton populations. The dynamics of both dissolved and particulate B-vitamin concentrations in space and time were striking - from subpicomolar to high picomolar levels observed and strong short-term (weeks) variability. We find notable autochtonous B-vitamin production in the estuary, but we expect the ability of the system to deliver these micronutrients to the ocean will depend on flushing as well as changes in microbial community. We identify vitamin B1, B12, psB12 (pseudocobalamin), and B3 as key explanatory variables for change in prokaryotic and eukaryotic NRE plankton, providing new evidence of B-vitamin influence upon estuarine plankton community composition. Our work reveals new complexities in B-vitamin production and consumption within zones of estuaries while underscoring these micronutrients as key drivers of microbial plankton composition.

## INTRODUCTION

Estuaries exhibit dynamic hydrological and physical-chemical properties which alter biology along the river to ocean continuum (Bianchi 2007). Microbial (prokaryotic and eukaryotic) communities drive biogeochemical cycling and primary production in aquatic environments – including estuaries – and the diversity and physiology of these microbes can vary notably across temporal and spatial scales (Peierls et al. 2012; Hall et al. 2013; Wang et al. 2020). Accordingly, identifying key factors that influence microbial communities over space and time is of significant interest. Monitoring inorganic (macro)nutrients in aquatic ecosystems and microbial community responses has been the focus of many research and monitoring efforts (Paerl 2006). For example, elevated nitrogen and phosphorus inputs are particularly impactful to estuarine microbial communities, leading to eutrophication and promotion of harmful algal blooms (HABs) (Wurtsbaugh et al. 2019).

The Neuse River Estuary (NRE, North Carolina) is a major tributary of the 2^nd^ largest estuary complex in the USA, the Albemarle Pamlico Estuarine System (APES), characterized by salinity and nutrient gradients, seasonality, and hydrological variation that impacts plankton abundance and composition along the estuary (Pinckney et al. 1998; Peierls et al. 2012; Hall et al. 2013; Gong et al. 2018). In the NRE, moderate to low river flow leads to biological heterogeneity – specifically a Chlorophyll *a* (Chl *a*) max mid-estuary and increased planktonic biomass. Similar to other temperate estuary systems, the main phytoplankton groups in NRE are chlorophytes, diatoms, dinoflagellates and cryptophytes (Pinckney et al. 1998; Gong et al. 2018, 2020) with high abundances of cyanobacteria during summer (Gaulke et al. 2010; Hall et al. 2013). Prokaryotic plankton community analyses in the NRE have focused on pathogens (e.g. *Vibrio* spp.) (Froelich et al. 2019) and recently cyanobacterial populations (Sánchez-Gallego et al. 2025), and dominant populations are expected to be similar to other temperate estuaries (Wang et al. 2020). Changes in inorganic nutrients and/or hydrological properties (e.g. salinity, discharge) explain some, but not all, of the observed variability in microbial plankton in the NRE (Peierls et al. 2012; Hall et al. 2013) and beyond. Micronutrients like tracemetals and B-vitamins are commonly overlooked and the impact of B-vitamins on microbial communities within estuary has not been robustly studied.

B-vitamins are a group of eight essential micronutrients that primarily function as enzyme cofactors in cells - but also as antioxidants or riboswitch ligands (Lukienko et al. 2000; Miranda-Ríos et al. 2001). Vitamin B1 (B1, thiamin), vitamin B2 (B2, riboflavin), vitamin B3 (B3, niacin or niacinamide), vitamin B5 (B5, pantothenate), vitamin B6 (B6, pyridoxine) and vitamin B12 (B12, cobalamin) are essential for most cells – with the exception of some organisms lacking B12 requiring enzymes altogether (e.g. SAR11 bacterioplankton) (Carini 2013). Most prokaryotic and eukaryotic plankton taxa are auxotrophic (unable to synthesize a vitamin they require for metabolism *de novo*, termed auxotrophs) for one or more B-vitamins (Tang et al. 2010; Sañudo-Wilhelmy et al. 2014; Paerl et al. 2018). As a result, these taxa require an exogenous source of the respective vitamin or vitamers (vitamin related compounds, such as precursors for *de novo* synthesis or degradation products) to meet cell needs.

Common NRE phytoplankton groups are likely to include auxotrophs and taxa with distinct vitamin requirements. Cyanobacteria are B1 prototrophs (organisms capable of *de novo* syntesis) and are hypothesized to be dominant summertime B1 synthesizers of coastal brackish environments (Sañudo-Wilhelmy et al. 2014; Bittner et al. 2024). Cyanobacteria also produce pseudocobalamin (psB12), a cobalamin analog that is not biologically available to most other plankton but can be made useful to other plankton groups after microbe-mediated chemical remodeling (Helliwell et al. 2016; Heal et al. 2017; Bannon et al. 2024b). We hypothesize that the availability of B-vitamins are important drivers of unresolved microbial plankton variability as seen in other marine systems (Paerl et al. 2018; Joglar et al. 2020; Bannon et al. 2025). There is also evidence of changes in microbial community composition during limiting B-vitamin conditions, potentially altering community composition of lower trophic levels and leading to vitamin deficiency at higher trophic levels (e.g. seabirds, fish) (Balk et al. 2009; Joglar et al. 2020). Therefore, quantification of B-vitamins and vitamers in aquatic ecosystems is of considerable interest as it potentially alters productivity, biomass, and/or the success of specific taxa. Only few studies simultaneously quantified multiple B-vitamins in the dissolved phase (Sañudo-Wilhelmy et al. 2012; Heal et al. 2014; Suffridge et al. 2017; Bruns et al. 2022, 2023; Bannon et al. 2025), and measurements of B-vitamins/vitamers in environmental plankton biomass are even more limited (Suffridge et al. 2017, 2018; Bannon et al. 2025).

Knowledge on B-vitamins and vitamers dynamics along the freshwater, brackish to coastal ocean gradient is lacking, and their connection to patchy biology are unknown - contrasting with well-described and modeled estuary macronutrient dynamics (Twomey et al. 2005). A few studies have quantified select B-vitamins/vitamers in a few brackish systems, and shown that their concentrations vary by two orders of magnitude between systems (Sañudo-Wilhelmy et al. 2006; Gobler et al. 2007; Koch et al. 2012; Heal et al. 2014; Tovar-Sánchez et al. 2016; Gómez-Consarnau et al. 2018; Bruns et al. 2022, 2023; Möller et al. 2022; Bittner et al. 2024). While some of this observed variability may be due to analytical difficulties and differences, overall, data on environmental concentrations of vitamins/vitamers are scarce and the effects of these micronutrients on microbial plankton growth and community dynamics in estuarine systems are not well understood. Thus, it remains unclear whether estuaries could be significant allochthonous sources of B-vitamins to the coastal ocean. Understanding these potential sources is important in a broader biogeochemical context, as the extremely low picomolar concentrations of dissolved B-vitamins in coastal and open ocean may limit microbial growth.

Here, we leverage substantial ongoing biogeochemical monitoring within the NRE and tackle the following research questions: (1) How do B-vitamin concentrations change short-term (weeks) along the salinity gradient of a temperate estuary (fresh to polyhaline)? (2) What are the relationships between B-vitamins and size-differentiated planktonic communities? To address these questions, we measured B-vitamins and vitamers in two particulate size fractions (picoplankton 0.22-3 µm; nano-/microplankton 3-90 µm) and in the dissolved phase using targeted liquid-chromatography mass spectrometry. Simultaneously, we characterized picoplankton (0.22-3 µm) and nano-/microplankton (3-90 µm) communities by 16S and 18S rRNA gene sequence analyses. We hypothesized that (1) B-vitamin concentrations are dynamic on short time scales and exhibit distinct patterns along the estuary relative to macronutrients; and (2) plankton community composition is significantly impacted by B-vitamin/vitamer availability, especially B1 and B12. The results provide a high resolution into the short-term dynamics of multiple B-vitamins and vitamers across phases and along a salinity gradient of a temperate estuary.

## MATERIALS AND METHODS

### Sampling

Near surface water (0.5 m) was collected on 26 July, 13 and 27 September, and 11 and 25 October 2021 from stations NRE0, NRE50, NRE70, NRE100, NRE120, NRE180 in collaboration with the University of North Carolina at Chapel Hill Institute of Marine Sciences (UNC-IMS) Neuse River Estuary Modeling and Monitoring Project (MODMON; https://paerllab.web.unc.edu/modmon/; Fig. 1A). NRE160 was sampled once on 11 Oct as weather conditions did not permit sampling at NRE180. Neuse River flow data was obtained from USGS gauge 02091814 near Fort Barnwell upstream of the NRE (USGS National Water Information System Web Interface: https://waterdata.usgs.gov/nwis; Supporting Information Fig. S1).

**Fig. 1.**
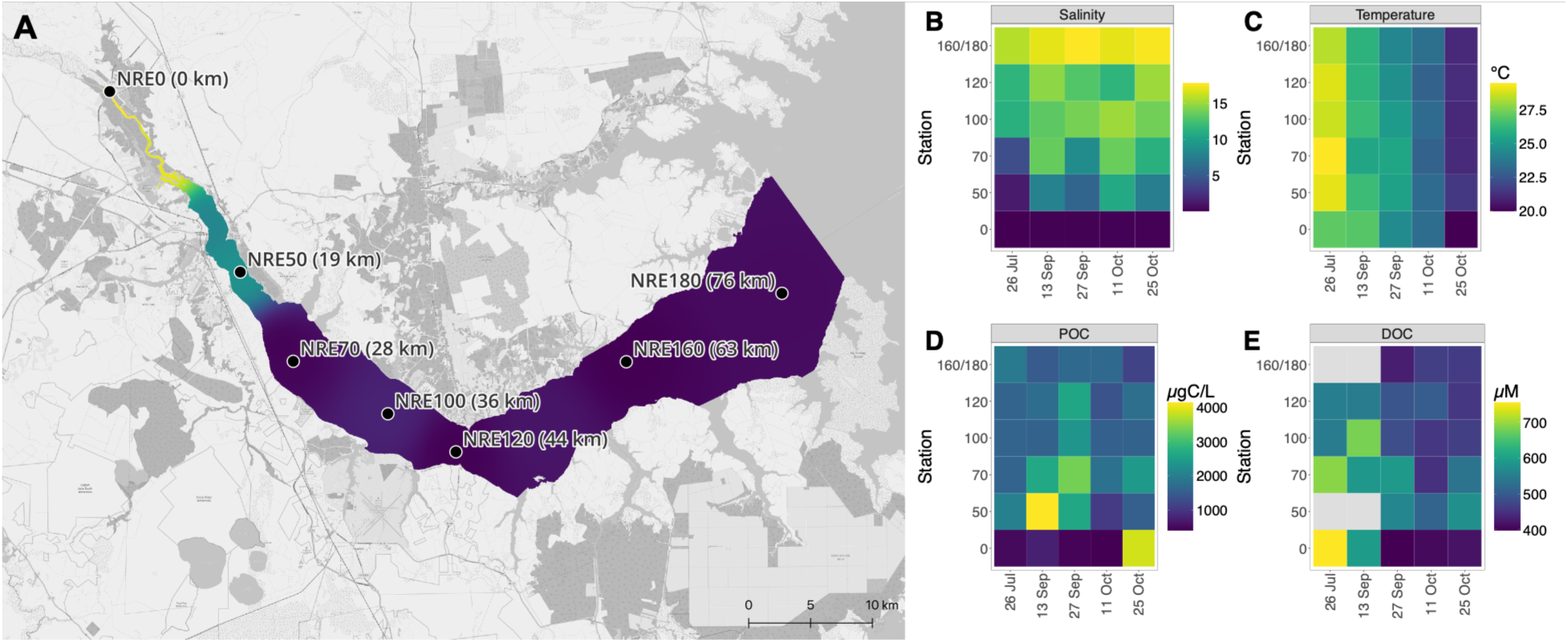
Environmental conditions at estuary sampling sites. Map of NRE stations sampled and distance along the river from station NRE0 is provided in parenthesis, and color gradient shows median N:P ratio along the estuary from the sampling time points (**A**), based on inorganic nutrient data from the MODMON monitoring program. Gradients in salinity (**B**), temperature (**C**), particulate organic carbon (POC, **D**) and dissolved organic carbon (DOC, **E**) across the stations and time points sampled. Data may be found in Supporting Information Data S1.

### Environmental parameters

Temperature, salinity, and turbidity were measured by YSI 6600 multi parameter quality sonde (Yellow Springs Instruments, Ohio, USA). Particulate Organic Carbon (POC) was measured by Costech ECS 4010 Analyzer as described before (Peierls et al. 2003). Dissolved Organic Carbon (DOC) was measured by Shimadzu TOC-5000A Analyzer as previously described (Crosswell et al. 2012). Extracted Chl *a* was determined fluorometrically with a Turner Trilogy Fluorometer (Peierls et al. 2012). Nitrate/nitrite (NO_3_^−^+NO_2_^−^), ammonium (NH_4_^+^), orthophosphate (PO_4_^3-^), total dissolved nitrogen (TDN) and silica (SiO_2_) were measured with a Lachat QuickChem 8000 Automated Ion Analyzer (Paerl et al. 2010; Peierls and Paerl 2010). Detection limits were 0.05 µM, 0.50 µM, 0.21 µM and 1.17 µM for NO_3_^−^+NO_2_^−^, NH_4_^+^, PO_4_^3-^ and SiO_2_, respectively. Primary productivity was assessed by light/dark ^14^C bicarbonate incorporation (Paerl 2006; Gaulke et al. 2010). These measurements are provided in Supporting Information Data S1.

### Bacterial and phytoplankton abundance

Whole water samples were kept in the dark on ice overnight and then fixed for flow cytometry counting the following day with glutaraldehyde (0.25% final) for 15 min in the dark at room temperature and stored at −80°C (Paerl et al. 2020). Prior to counting, samples were thawed until they reached room temperature. Bacterioplankton and phytoplankton abundances were determined using a Guava EasyCyte HT (Millipore) flow cytometer equipped with red and blue excitation lasers. Phytoplankton were counted based on fluorescence (Paerl et al. 2020) with additional gating based on forward scatter to enumerate single cells rather than aggregates (Supporting Information Fig. S2). Bacterioplankton were counted using SYBR Green I staining and without any heating step (Brussaard et al. 2010). Abundances are provided in Supporting Information Data S1.

### Metabolite sample collection and analysis

Sampling bottles, filtration units, and collection bottles (amber, HDPE) were cleaned with 0.1 M HCl, methanol, and MilliQ water. Sampling bottles were rinsed with sample water, filled with water through a 90 µm Nitex mesh, and stored at near in-situ temperature in the dark. Water was filtered within 10 h in a dark room through 3.0 µm polycarbonate (Isopore, Millipore) and 0.20 µm nylon filters (Millipore). The biomass from the nano- and microplankton fraction that passed through the 90 µm mesh and was retained on the 3.0 µm filters was for simplicity defined as “microplankton”, and the biomass fraction that was sequentially retained on the 0.2 µm filter was defined as “picoplankton”. Three bottles of ∼250 mL filtrate per station were prepared and stored at −20°C.

Metabolite extractions were conducted in a dark room with a red headlamp as light source. Dissolved metabolites were captured using C_18_ Solid Phase Extraction (SPE; waters, 10 g) columns and triplicate extractions were performed for each sample. Filtrates were thawed at 4°C, pH adjusted to 6.5 and spiked with ^13^C-thiamin (thiamine-4-methyl-^13^C-thiazol-5-yl-^13^C_3_, Sigma-Aldrich, 75 pM final). Particulate metabolites were extracted from 3.0 and 0.20 µm pore-size filters (Heal et al. 2014), and prior to extraction, vials with sample filters were spiked with 10 pmol ^13^C-B1 (4,5,4-methyl-^13^C_3_, 97%; Cambridge Isotope Laboratories), 1 pmol heavy B2 (^13^C_4_-^15^N_2_, 97%; Cambridge Isotope Laboratories), and 2 pmol cyano-cobalamin (Fisher BioReagents). Mean percent recoveries of ^13^C-B1 are provided in Supporting Information Table S1. Eluted metabolites were analyzed using a Dionex Ultimate-3000 LC system coupled to a TSQ Quantiva triple-stage quadrupole mass spectrometer (ThermoFisher) operated in selected reaction monitoring mode. Matrix groups included a high salinity grouping (NRE180, NRE160, NRE120, NRE100) and a low salinity grouping (NRE70, NRE50, NRE00), as matrix differences were expected. For each matrix group limits of detection (LOD) and limits of quantification (LOQ) were determined by calculating (x3) and (x10) the standard deviation of the QC pool run between samples, respectively (Supporting Information Data S2). For further details see (Paerl et al. 2023a) or additional Methods in the Supporting Information.

Data analysis was adapted from (Heal et al. 2014; Bannon et al. 2025). For particulate samples, the peaks of B1 and B2 were normalized to the stable isotope internal standard to reduce variability from the instrument and sample preparation. Metabolite measurements were excluded if only one of the two analytical injections was above the LOD. The mean of the two analytical injections was calculated before applying percent recoveries. Particulate B1 was corrected for percent recovery of the stable isotope internal standard as variable recovery was observed (Supporting Information Table S1). Dissolved B1 was corrected for percent recovery of the ^13^C-B1 spike in each sample. Dissolved HMP, HET, FAMP and cHET were corrected for percent recovery previously determined from samples from the estuary (Paerl et al. 2023a), for matrix groups high and low salinity recovery values from station NRE180 and station NRE0 were applied, respectively (Supporting Information Table S1). The mean and standard deviation were calculated from biological replicate samples that passed LOD and LOQ, or batch by batch criteria if between LOD and LOQ (Supporting Information Data S3). A few samples were lost during sample processing or excluded during analysis see Supporting Information Methods.

Metabolite data was analyzed and visualized with heatmaps (R package ComplexHeatmap v2.20.0), LODs and LOQs were included with their calculated picomolar concentration. Metabolite concentrations (mean of biological replicates) are shown relative to the concentration range of each compound (rows). Each sample (columns) is made up of the different relative vitamin concentrations, referred to here as a vitamin profile, like a B-vitamin “fingerprint” from that sample. Columns and rows were clustered based on Euclidean distances corresponding to differences between samples and relative metabolite concentrations, respectively. To analyze the phase partitioning of vitamins, only measurements above LOD/LOQ were included and visualized.

### DNA extraction and sequencing

Sampling bottles were rinsed with sample water, filled with water through a 90 µm Nitex mesh and stored at near *in-situ* temperature in the dark. Within six hours of sampling, 1 L of water was sequentially filtered onto a 3.0 µm pore-size membrane (MCE, Millipore) and a 0.22 µm pore-size Sterivex filter (PES, Millipore), which were stored at −20°C. DNA was extracted with the DNeasy Blood and Tissue kit (Qiagen) with additional lysozyme and proteinase K steps (Bittner et al. 2024) and quantified (Qubit 3.0, Invitrogen).

Partial 16S and 18S rRNA genes were PCR amplified from both size fractions (0.22-3.0 µm, 3-90 µm) with the KAPA HiFi HotStart ReadyMix (Roche) and primer pairs 515F-Y/926R (Parada et al. 2016) and a modified version of 565F/964R primers to avoid mismatches with haptophytes (Lin et al. 2017), previously used for NRE plankton community analysis (Gong et al. 2020). Primer sequences and PCR conditions are provided in Supporting Information Table S2. Triplicate PCR reactions were pooled, amplicons were indexed and sequenced with MiSeq 2 x 300 bp v3 (Illumina) at the Rush Genomics and Microbiome Core Facility, Rush University, Chicago, IL, USA (Naqib Ankurand Poggi 2018).

### Sequence and data analysis

Reads were trimmed with cutadapt (v3.4) (Martin 2011), quality filtered (16S: F 277, R 240; 18S: F 280, R 250), dereplicated, denoised, read pairs were merged (minOverlap = 100) and chimeras removed in DADA2 (v1.22.0) (Callahan et al. 2016). For both 16S and 18S rRNA gene fragments, Amplicon Sequence Variants (ASVs) were generated. Taxonomy was assigned to 16S ASVs with ‘assignTaxonomy’ from DADA2 with the SBDI-curated version (v7) (Lundin and Andersson 2021) of 16S sequences of GTDB (r09-rs220) (Parks et al. 2018; Pascoal et al. 2024). Taxonomy to 18S ASVs was assigned with ‘IdTaxa’ from DECIPHER (Murali et al. 2018) with the PR2 (v5.0.0) database (Guillou et al. 2012). Relative abundances of dinoflagellates may be overestimated due to their high 18S rRNA copy number relative to diatoms (Gong and Marchetti 2019).

Both 16S and 18S ASV tables were rarefied (n = 100) to the lowest sequencing depth (16S: 15,031 reads; 18S: 1,490 reads) with the ‘rrarefy’ function from the vegan package (v2.6.8) (Oksanen et al. 2022), to account for varying sequencing depth in accordance with recent recommendations (Schloss 2024). One sample from a 3.0 µm filter from station NRE180 from 25 Oct was removed due to insufficient sequencing depth prior to rarefaction. Further processing was carried out in phyloseq (v1.46.0) (McMurdie and Holmes 2013). Sequences without a domain annotation and singletons were removed. Empty taxonomic levels were filled by the nearest classified taxonomic level with ‘tax_fix’ from microViz (v0.12.10) (Barnett et al. 2021).

ASV abundance data was Hellinger transformed, environmental and metabolite data was z-score transformed prior to transformation-based Redundancy Analysis (tb-RDA) and R2-adjustment with vegan. Explanatory variables for a constrained ordination were selected by forward selection by permutation (nperm = 999) of residuals under reduced model by ‘forward.sel’ implemented in adespatial (Dray et al. 2012). Significance of the constrained tb-RDA, the axes and the explanatory variables were tested with ‘anova.caa’ from the vegan package.

Principal Coordinate Analyses (PCoA) were performed with vegan on Bray-Curtis distance matrixes of Hellinger transformed abundance data. Correlation analyses were conducted with rstatix (v0.7.2, A. Kassambara: https://github.com/kassambara/rstatix) and visualized with corrplot (v0.92) (Wei and Simko 2021), the lower LOD (Supporting Information Data S1) was used for measurements below LOD/LOQ. Linear models were fitted with the ‘lm’ function implemented in R. Data was analyzed and visualized in the R environment (v4.4.1) with tidyverse (v2.0.0) (Wickham et al. 2019).

## RESULTS

### Hydrological and biochemical conditions in the estuary

NRE surface water salinity ranged from 0 to 17 – peaking at station NRE180 furthest down the estuary (Fig. 1B). During high river discharge (Supporting Information Fig. S1A) a notable freshwater signature occurred further downstream, e.g. periods of salinity ∼4 at NRE70 (Fig. 1B). Discharge (Neuse River flow) was elevated between the samplings of 11 and 25 October (Supporting Information Fig. S1A). Surface water temperature peaked at 29°C in July and declined to 21°C during October (Fig. 1C), showing a seasonal shift from summer to fall. NOx concentrations were below detection limit (< 0.05 µM), except at NRE0 where concentrations reached up to 57 µM on 13 September. Ammonium was mostly below detection limit (< 0.50 µM) but was elevated on 11 October (2.3-11.1 µM). Phosphate concentrations ranged from < 0.21 to 2.9 µM and showed higher levels on 13 September. The ratio of nitrogen to phosphate was typically highest (>20) at NRE0 and decreased towards the middle of the estuary (NRE70; Fig. 1 A), with the execption of 25 October. Between 11 and 25 October, inorganic nutrients became depleted at all stations after a period of higher discharge, change was especially pronounced at NRE0 (Supporting Infromation Data S1). Silica concentrations followed the salinity gradient, with higher concentrations upstream (Supporting Information Fig. S3B). Dissolved organic nitrogen (DON) averaged 292 µg N L^−1^ with periodically higher concentrations at NRE0 (e.g. 26 Jul 420 µg N L^−1^). Dissolved organic carbon (DOC) ranged from 399 to 754 µM, both minimum and maximum were observed at NRE0 (Fig. 1E). As is typical for the NRE (Gaulke et al. 2010), maximum Chl *a* occurred mid-estuary (NRE50, 70) reaching 20-40 µg L^−1^ (Supporting Information Fig. S3D). A notable exception occurred on October 25th, where 90 µg Chl *a* L^−1^ occurred in NRE0 surface water coinciding with peaks in turbidity (13.1 NTU), Particulate Organic Carbon (POC, 3.9 mg C L^−1^, Fig. 1D) and primary productivity (277 mg C m^−3^ h^−1^; Supporting Information Fig. S3).

### Quantification of dissolved B-vitamins and vitamers

Twelve B-vitamins and vitamers were quantified from 22 dissolved phase sampling events (60 samples; full list found in Supporting Information Data S3A, Fig. S4). The biologically active forms of B12 (Ado-B12, Me-B12) and psB12 (Me-psB12) were not detected, as expected due to in-situ photodegradation into OH-B12 and OH-psB12, respectively (Bannon et al. 2025). Dissolved OH-B12 ranged from 0.9 pM up to 3.0 pM (Fig. 2A), and OH-psB12 concentrations were notably lower - approximately half (0.4 −1.4 pM) that of OH-B12 and occasionally below our detection limit (0.3 pM). Concentrations of DMB, the alpha ligand of B12, were above the LOD of 2.8 pM but not quantifiable in most samples. Temporally and spatially, concentrations of dissolved B1 ranged from 19 to 74 pM, while concentrations of B1 vitamers (HMP, AmMP, FAMP, cHET, HET) were lower and ranged from (below limit of detection - cHET, HMP, AmMP; Supporting Information Data S2A) to 82 pM for FAMP – a B1 degradation product.

**Fig. 2.**
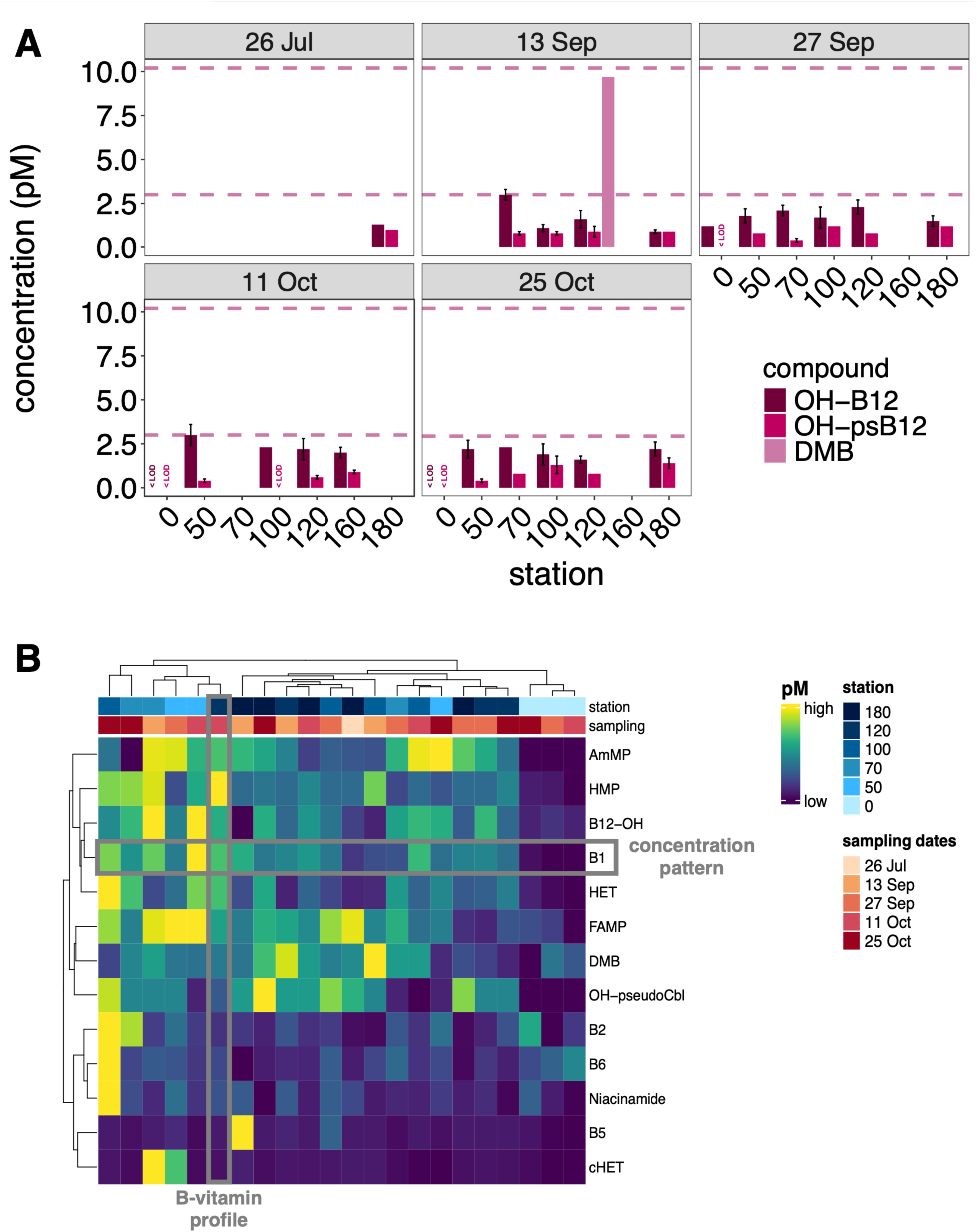
Dissolved B-vitamins and vitamers. Bar plot showing concentrations of dissolved OH-B12, OH-psB12 and B12 lower ligand DMB (**A**). Upper and lower dashed lines indicate the average LOQ and LOD for DMB, respectively. The average LOD and LOQ for OH-B12 was 0.9 pM and 3.4 pM, respectively. The average LOD and LOQ for OH-psB12 was 0.3 pM and 1.3 pM, respectively. Samples below LOD are indicated; error bars show ± standard deviation of biological replicate water samples. Heat map of dissolved B-vitamins and vitamer profiles (**B**). Rows represent concentration patterns across samples for each compound and columns represent B-vitamin profiles for each station and sampling date (illustrated by the gray boxes). The mean metabolite concentrations across water replicates are shown relative to the concentration range of each compound; clustering is based on Euclidean distances. For measurements below LOD or LOQ, the concentration limits are displayed (Supporting Information Data S2A).

Dissolved B2 concentrations (12-96 pM) were comparable to B1 but with a notable increase across the estuary on Oct 25 following high discharge. Similary to B2, B6 was detected in all samples and ranged from 7-36 pM (mean 13.7 ± 6 pM). B5 was notably low (< LOD) at higher salinity stations (NRE100, 120, 180), with a unique peak of 159 ± 21 pM at NRE180 on 13 September. With respect to variability, B1 pyrimidine vitamers (HMP, AmMP, FAMP) showed moderate variation (2 to 4-fold change) while B5 (159 ± 21 pM peak; 17 ± 14 pM mean; 13 Sep NRE180) and cHET (628 ± 153 pM peak; 58 ± 13 pM mean, 27 Sep NRE50) concentrations were stable, except occasional high peaks (10-fold higher; Fig. 2B). Concentration patterns of dissolved B1 and OH-B12 clustered together (clustering of rows), but relative concentrations of dissolved OH-psB12 concentrations showed a unique pattern compared to the other vitamins with higher concentrations (∼4-fold) in the lower estuary (NRE100, 120, 180; Fig. 2). Concentration patterns of dissolved B2, B6 and B3 clustered together and were characterized by a maximum at NRE100 on 25 Oct. This signature of increased dissolved B2 was also detected upstream at NRE70 on 25 Oct.

Freshwater dissolved vitamin samples (NRE0) clustered together and showed overall lower relative concentrations compared to downstream stations (Fig. 2B). Interestingly, higher-salinity samples from NRE180 clustered with the freshwater station cluster and most NRE120 samples, whereas brackish mid-estuary stations (NRE50, NRE70) formed a separate cluster with some NRE100 and NRE120 samples – jointly pointing to the middle estuary as site of unique dissolved vitamin/vitamers composition.

### Quantification of particulate B-vitamins and vitamers

Sixteen B-vitamin and vitamers were detected from 23 particulate sampling events in two size fractions (0.2-3 µm - 90 samples, 3-90 µm - 58 samples; Fig. 3; full list found in Supporting Information Data S3B, C). Mean particulate B1 concentration (0.2-3 µm: 25 ± 18 pM, 3-90 µm: 21 ± 16 pM) and its range (LOD to ∼70 pM) were similar for both size fractions. HET in the 0.2-3 µm size fraction was only quantifiable at five stations (NRE0, 50, 70, 100, 120) on 25 Oct, following higher discharge, with a maximum of 3.6 ± 0.6 pM at NRE120. Peak HET concentration occurred in the 3-90 µm size fraction on 25 Oct at NRE120 and NRE0 reaching ∼12.5 pM, otherwise concentrations were below ∼1-2 pM. Traces of cHET were in both size fractions but not quantifiable due to low signal to noise ratios and low concentrations. FAMP concentration was similar in both particulate size fractions (0.2-3 µm: 2.8-18.1 pM, 3-90 µm: 4.3-20.2 pM). In contrast, HMP was only quantifiable in select samples from the 3-90 µm size fraction from the 13 and 27 Sep with concentrations of up to 14 ± 7 pM at NRE120. Contrasting with dissolved concentrations, AmMP reached upwards of 65 ± 10 pM (3-90 µm) at NRE120 and NRE180 on 25 Oct with notable variation between samples (non-quantifiable in other samples; Supporting Information Data S3).

**Fig. 3.**
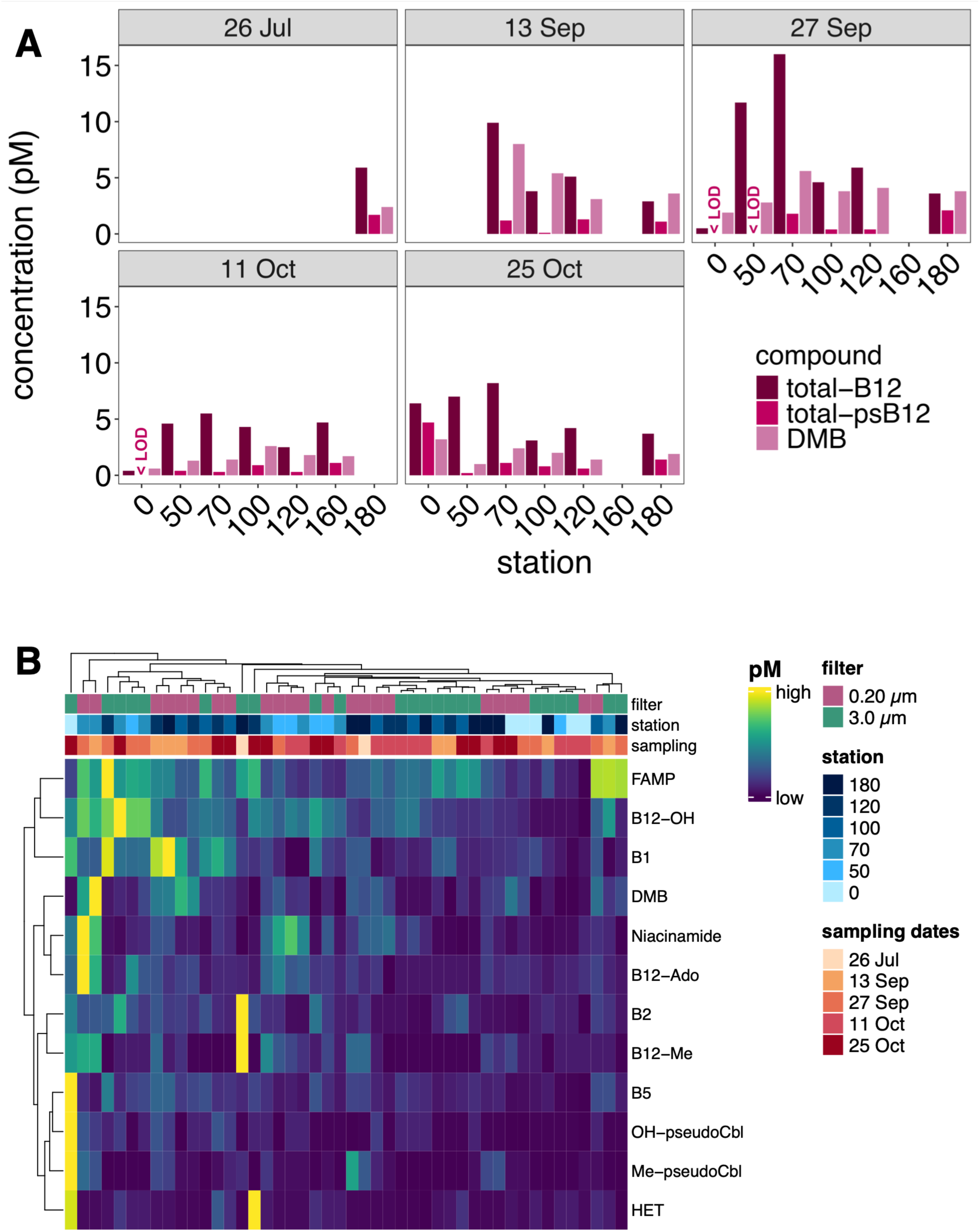
Particulate B-vitamins and vitamers. Bar plot showing concentrations of particulate (0.2-90 µm) total-B12, total-psB12 and B12 lower ligand DMB (**A**). Samples where total-psB12 was below the limit of detection of 0.1 pM is indicated; error bars show ± standard deviation of biological replicate water samples. Heat map of size-fractionated particulate B-vitamins and vitamer profiles (**B**). Rows represent concentration patterns across samples for each compound and columns represent B-vitamin profiles for each station and sampling date. The mean metabolite concentrations across biomass replicates are shown relative to the concentration range of each compound; clustering is based on Euclidean distances. For measurements below LOD or LOQ, the concentration limits are displayed (Supporting Information Data S2B, C).

Mean particulate B2 varied two-fold between size fractions (0.2-3 µm: 4.3 ± 2.9 pM, 3-90 µm: 9.0 ± 8.2 pM) with lowest concentrations (< 2 pM) across the estuary occurring 11 Oct. B3 in the 0.2-3 µm size fraction was highly dynamic with variations of more than 20-fold ranging from 14 pM to a of maximum concentration of 306 ± 42 pM at NR70 on 27 Sep. In contrast, B3 in the 3-90 µm size fraction were less dynamic with a mean of 20 ± 17 pM, excluding a peak concentration of 125 pM at NRE0 on 25 Oct. Generally, particulate B5 in both size fractions was lowest at freshest station NRE0 (< 2 pM), except on 25 Oct where a maximum (50 ± 19 pM in the 3-90 µm size fraction) was found at NRE0.

Three forms of B12 (Me-B12: methylcobalamin, Ado-B12: adenosylcobalamin, OH-B12: hydroxycobalamin) and two forms of psB12 (OH-psB12: hydroxy-pseudocobalamin, Me-psB12: methyl-pseudocobalamin) could be resolved in the particulate size-fraction, but bioavailable forms (Ado-B12, Me-B12, Me-psB12) were often below their respective LODs, especially in the 3-90 µm size fraction. Mean OH-B12 concentration was 1.1 ± 0.5 pM and 1.3 ± 0.8 pM in 0.2-3 µm and 3-90 µm size fractions, respectively. The maximum particulate concentration for any B12-compound was Ado-B12 at NRE70 on 27 Sep with 7.6 ± 2.8 pM in the 0.2-3 µm size fraction, highlighting a biological active form of B12 can be twice the concentration of OH-B12. Concentrations of DMB (mean 2.2 ± 1.4 pM) were patchier than B12 forms in the 0.2-3 µm size fraction and between the LOD and 1.7 pM in 3-90 µm size fraction (Fig. 3A, Supporting Information Fig. S5). Notably, on 25 Oct 2021 3.4 ± 0.6 pM Me-psB12 was detected in the microplankton.

Clustering based on Euclidean distances was performed to identify similarities and differences across particulate metabolites, stations and sampling dates (Fig. 3B). FAMP, OH-B12, and B1 exhibited similar variability in concentration patterns of particulate vitamin concentrations (relative to the concentration range of the respective compound) across sampling dates, stations, and the two size fractions (Fig. 3B; clustering of rows in heatmap). These three compounds (FAMP, OH-B12, B1) were characterized by occasionally elevated concentrations, contrasting to other compounds (e.g. B2), which had sporadic sometimes 10-fold higher concentrations. Particulate vitamin profiles from the two size fractions did not exhibit clustering by station, sampling time, or size fraction (Fig. 3B). Only in four instances did vitamin profiles from the two size fractions at the same station and time cluster closely together (e.g., NRE0 on 27 Sep and 11 Oct, NRE120 on 11 Oct, NRE70 on 25 Oct). Overall, particulate vitamin profiles from a given size fraction were more similar to profiles from the same size fraction at other stations than to the alternate size fraction at the same station (e.g. 0.2-3 µm profiles from NRE100 and NRE120 on 11 Oct).

When clustering the particulate vitamin profiles per size-fraction separately, the three freshwater (NRE0) vitamin profiles cluster together in the 0.2-3 µm size fraction as they were characterized by overall lower concentrations contrasting to samples from the middle estuary (NRE70, 100) (Supporting Information Fig. S6). Highest particulate vitamin concentrations in the 0.2-3 µm size fraction were measured at NRE70 in September. The particulate vitamin profile of the 3-90 µm size fraction at NRE0 from 25 Oct, was unique with peak concentrations of B5, psB12 and HET grouping separately (Fig. 3).

### B-vitamins and abiotic estuary conditions

A Kendall’s rank correlation analysis was performed for metabolites with each other and a suite of associated measurements (Fig. 4A, Supporting Information Fig. S7). To investigate if rivers could be a source of vitamins to coastal systems, B-vitamin and vitamer concentrations were analyzed for correlations with salinity, and results were vitamin-specific (Fig. 4). Dissolved B2 and B6 were negatively correlated with salinity. Dissolved OH-psB12 was positively correlated with salinity and higher concentrations aligned with higher abundances of picocyanobacteria, producers of pseudocobalamin (Bannon et al. 2024b) (Fig. 4B, Supporting Information Fig. S8). In the 0.2-3 µm particulate size fraction B5 and Me-psB12 were positively correlated to salinity. B1 and B3 within the 3-90 µm particulate size fraction were negatively correlated with salinity.

**Fig. 4.**
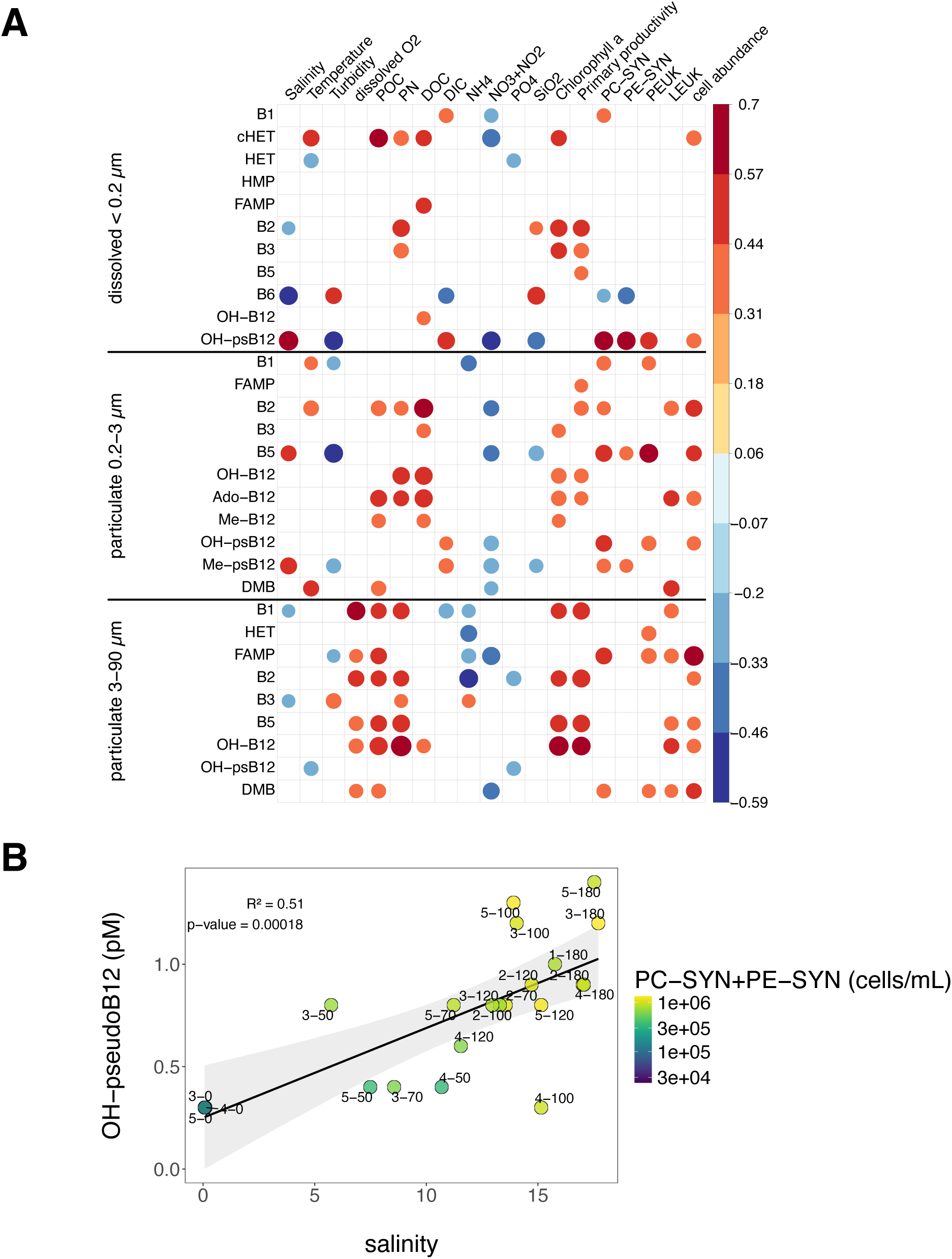
Correlation matrix of significant (p < 0.05) Kendall’s rank correlations between metabolites and environmental, nutrient and biological measurements (**A**). Linear relationship between dissolved OH-psB12 and salinity and corresponding cell abundances of *Synechococcus*-like phytoplankton (**B**). Numbers next to points indicate sampling and station number. Particulate organic carbon (POC), particulate nitrogen (PN), dissolved organic carbon (DOC), dissolved inorganic carbon (DIC) are abbreviated. Abundances of *Synechococcus*-like phycocyanin (PC)-rich cells (PC-SYN), *Synechococcus*-like phycoerythrin (PE)-rich cells (PE-SYN), picoeukaryotic phytoplankton cells (PEUK), larger eukaryotic phytoplankton cells (LEUK) and bacterial cell abundance (bacterial abundance) were determined by flow cytometry. For abbreviation of metabolites names see Supporting Data S2. Metabolites below LOD or LOQ were replaced by the lower LOD value prior to correlation analysis. Gray shading indicates the 95% confidence interval around the linear regression.

Next, we examined if patterns in B-vitamins/vitamers were connected to patterns of measured dissolved and particulate nutrients (Fig. 4A). Multiple dissolved and particulate vitamins (e.g. B1) displayed negative correlations to dissolved inorganic nitrogen forms, whereas multiple particulate B-vitamin concentrations were positively correlated to POC and DOC concentrations. Particulate B-vitamins (B2, B3 and B12) of the 0.2-3 µm size fraction were positively correlated to DOC concentrations. B2 was positively correlated to particulate nitrogen and Chl *a* in the dissolved and both particulate phases. Particulate B-vitamins (B1, B2, B3, B5, OH-B12) from the 3-90 µm size-fraction showed strong positive correlations to Particulate Nitrogen (PN) concentrations.

### Partitioning of B-vitamins across dissolved and particulate phases

To investigate vitamin connectivity between the dissolved and particulate pool, the concentrations of vitamins from both particulate size fractions were summed. Ratios of particulate to dissolved B1, B3 and B5 varied over time and fell on both sides of the identity line, a phase partitioning of 1:1 (Fig. 5A). Particulate total B12 concentrations were higher than dissolved OH-B12 concentrations in most samples, whereas concentrations of B2, FAMP and HET were higher in the dissolved pool than the particulate pool. When particulate vitamin/vitamer concentrations were normalized to POC and dissolved concentrations to DOC, patterns shifted towards higher values in the particulate size fraction (Supporting Information Fig. S9).

**Fig. 5.**
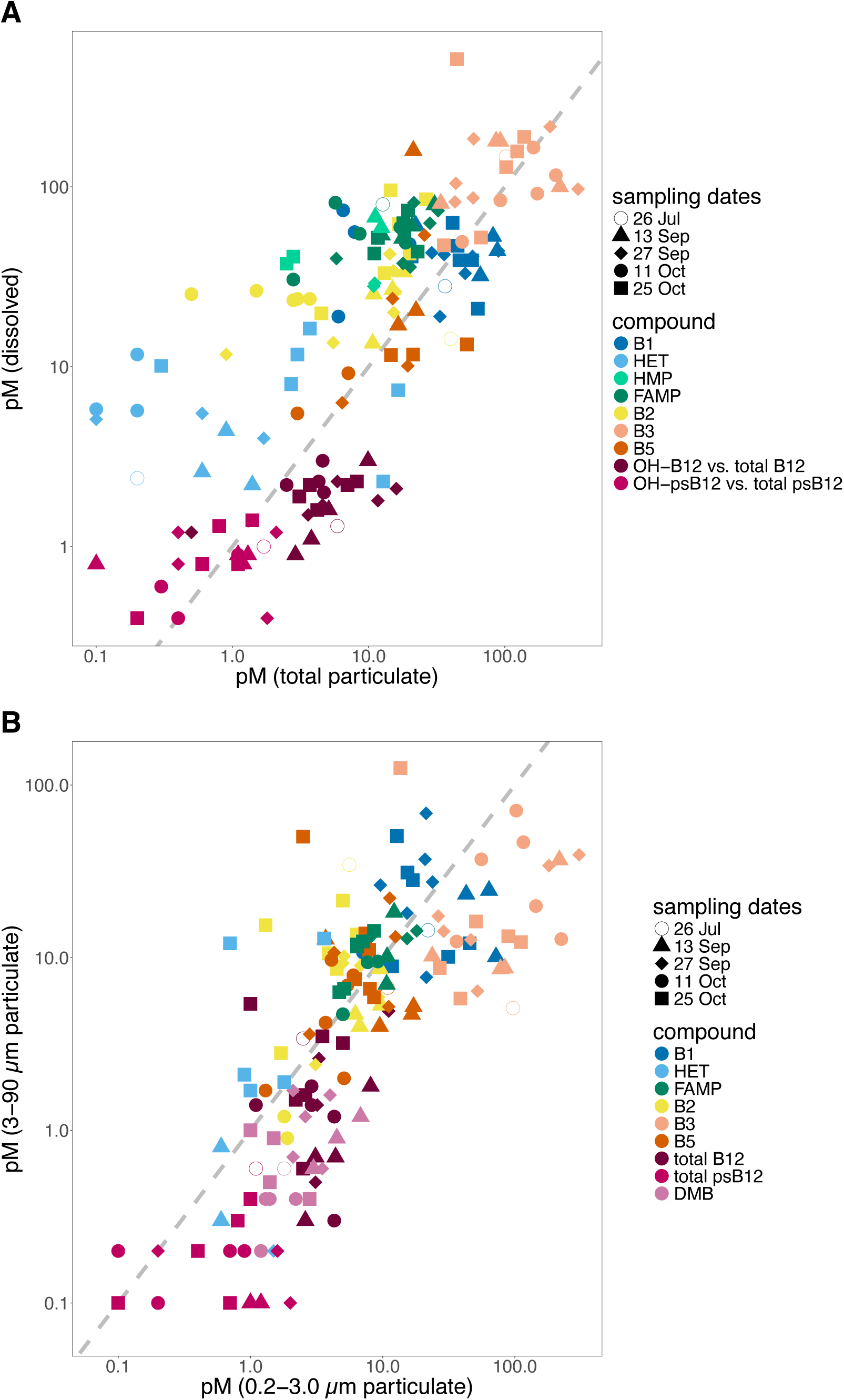
Scatter plots of phase partitioning of B-vitamins and vitamers, dissolved (< 0.2 µm) vs. particulate (0.2-90 µM) (**A**), particulate (3-90 µm) vs. particulate (0.2-3.0 µm; **B**). Axes are log10 scale and show picomolar concentrations. Shape of points indicates sampling time point and color corresponds to metabolite compound. Gray dashed line indicates a phase partitioning of 1:1. Points left of the line indicate an enrichment in the phase of the y-axis while points right of the line indicate an enrichment in the phase on the x-axis. Only metabolites with measurements in both phases were included; measurements below LODs were excluded. Particulate total B12 is the sum of Ado-B12, Me-B12 and OH-B12. Particulate total psB12 is the sum of OH-psB12 and Me-psB12.

When comparing particulate B-vitamin and vitamer concentrations between the two size fractions, B1, B2, B5 and FAMP showed temporal dynamics, but overall measurements fell around the identity line, where the concentration of a compound would be equal in both size fractions (Fig. 5B). Distinctly, total-B12, DMB and B3 concentrations in the 0.2-3 µm size fraction were higher compared to the 3-90 µm size fraction, except B3 on 25 Oct at NRE0. HET concentrations were often elevated in the 3-90 µm size fraction, whereas OH-psB12 were typically higher in the 0.2-3 µm size fraction.

B1 was the only compound exhibiting a spatial pattern across the two particulate pools (Supporting Data 3B, C). Particulate B1 concentrations were spatially distinct with higher concentrations (31 ± 21 pM) in the lower estuary (NRE180, 120, 100) in the 0.2-3 µm size fraction compared to the 3-90 µm size fraction (18 ± 17 pM); however, in the upper estuary (NRE0, 50, 70) B1 was higher in the 3-90 µm fraction (25 ± 13 pM). The most notable temporal feature observed was a sharp increase in B2, B3, B5, and HET concentrations in the 3–90 µm size fraction compared to the 0.2–3 µm fraction at NRE0 on 25 Oct Oct relative to the other time points sampled.

As an indicator of potential connections between metabolite pools, we analyzed correlations between quantified metabolites with Kendall’s rank correlation analysis (Supporting Information Fig. S7). Dissolved B1 was positively correlated with HET, while dissolved B3 positively correlated with dissolved B5. Most dissolved B-vitamins and vitamers were not correlated to their particulate concentration of either size fraction. In the 0.2-3 µm particulate size fraction some metabolites were positively correlated with each other (e.g. B3 and OH-B12) and metabolites of the two particulate size fractions exhibited only few significant (p < 0.05) correlations with each other.

### Plankton abundances

Four small phytoplankton morphotypes were detected in NRE surface water based on flow cytometry (Supporting Information Fig. S2, S3E-I). *Synechococcus*-like phycoerythrin (PE)-rich (PE-SYN) cells ranged from 7.29 × 10^1^ to 1.23 × 10^5^ cells mL^−1^, while *Synechococcus*-like phycocyanin (PC)-rich cells (PC-SYN) were more abundant (2.31 × 10^4^ to 1.62 × 10^6^ cells mL^−1^). PE-SYN was more abundant at higher salinity stations and peaked at NRE180, whereas PC-SYN cells were abundant throughout the estuary, except NRE0. Picoeukaryotic phytoplankton (PEUK) surged at NRE100 on 25 October, reaching 1.58 × 10^5^ cells mL^−1^. Abundance of larger eukaryotic phytoplankton cells (LEUK, Supporting Information Fig. S2; example plot with gates) peaked at NRE70 on 27 September with 2.66 × 10^5^ cells mL^−1^. Bacterioplankton abundance was on average 1.52 × 10^7^ cells mL^−1^ across stations and samples, except NRE0, where cell abundances were typically 10 x lower, with 4.24 × 10^6^ cells mL^−1^.

### Eukaryotic plankton community composition

A total of 1,640 eukaryotic ASVs were retained for analysis, with 19% unique to the microplankton size fraction and 32% unique to the picoplankton size fraction. The major eukaryotic plankton divisions present were *Stramenopiles*, *Alveolata, Cryptophyta* and *Chlorophyta* (Fig. 6A, Supporting Information Fig. S8). *Stramenopiles* were abundant in both size fractions (0.2-3 µm: 22 ± 15% relative abundance, 3-90 µm: 34 ± 19% relative abundance). The diatom genus *Cyclotella* was highly abundant, especially in September, accounting for up to 53% and 70% of relative abundance in the pico- and microplankton, respectively (Supporting Information Fig. S10), in line with peak cell abundances of the large eukaryote phytoplankton morphotype measured by flow cytometry (up to 2.7 × 10^5^ cells/mL; Supporting Information Fig. S3E). The *Alveolata* subdivision *Dinoflagellata* showed higher relative abundances in the microplankton size fraction (0.2-3 µm: 11 ± 11%, 3-90 µm: 26 ± 18%) and contributed up to 63% in relative abundance, whereas *Ciliophora* showed similar relative abundances in the two size fractions (0.2-3 µm: 10 ± 9%, 3-90 µm: 9 ± 10%) with peak relative abundances at NRE0 (mean 25 ± 11%). The *Dinoflagellata* genera *Polykrikos*, *Levanderina* and *Gyrodinium* were among the top 10 assigned genera (48% of taxa were not assigned a genus) based on relative abundance across the dataset and occurred predominantly in the microplankton. Peak relative abundances for *Polykrikos*, *Levanderina* and *Gyrodinium* were 42% (NRE180), 32% (NRE70) and 20% (NRE120) on 26 July, respectively (Supporting Information Fig. S9).

**Fig. 6.**
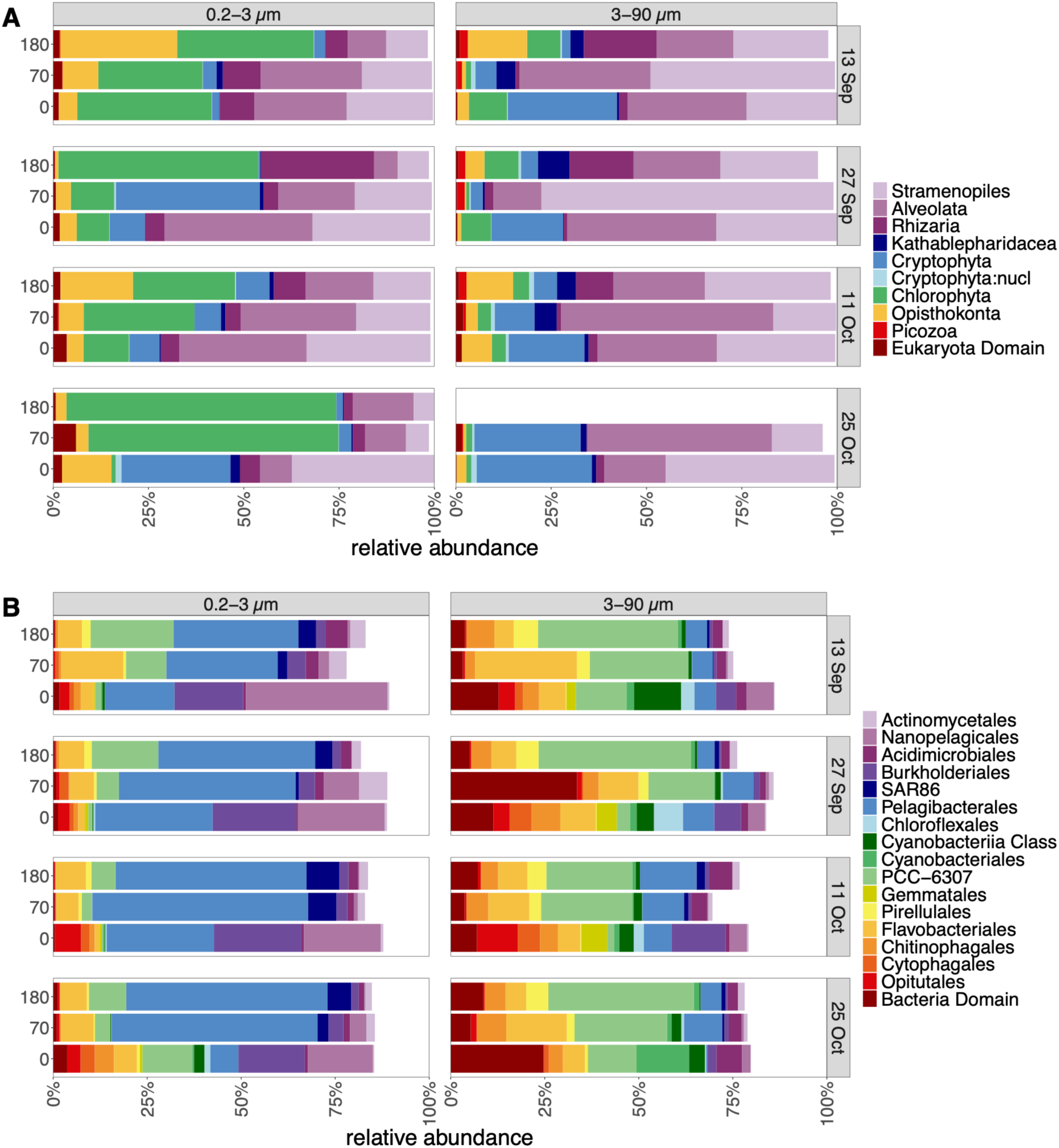
Eukaryotic (**A**) and prokaryotic (**B**) plankton community composition based on 18S and 16S rRNA genes at stations NRE0, NRE70 and NRE160/180. Taxonomy shown on division and order level for 18 and 16S rRNA, respectively. The sample NRE180 from the microplankton size fraction was removed due to low sequencing depth for the 18S rRNA gene amplicons. Taxonomic profiles for all sampled stations are provided in Supporting Information Fig. S10 and S13.

The dominant genus of *Cryptophyta* was *Cryptomonas* and occurred across stations and both size fractions but showed highest relative abundances at stations NRE50 and NRE70 on 27 September with 42 and 35% relative abundance, respectively (Supporting Information Fig. S11). *Chlorophyta* were present in higher relative abundances in the picoplankton (33 ± 17%) than in the microplankton (6 ± 3%; Fig. 6A). The genera *Bathycoccus* (e.g. 57% at NRE180 25 Oct) and *Micromonas* showed higher relative abundances in the lower estuary compared to *Ostreococcus*, which peaked in the upper estuary (e.g. 46% at NRE50 25 Oct, Supporting Information Fig. S11). Spikes in *Fungi* occurred in single samples (picoplankton size fraction), e.g. *Opisthokonta* (21% relative abundance) at NRE100 on 11 October and *Rhizophydiales* (13% relative abundance) at NRE0 on 25 October.

### Prokaryotic plankton community composition

In total, 4,451 prokaryotic ASVs were detected, 47% were unique to the microplankton size fraction, and 15% were unique to the picoplankton size fraction. Overall, a high number of 1,484 ASVs (33%) could not be classified beyond the order level. Within the prokaryotic picoplankton (0.2-3 µm) the bacterial phyla *Pseudomonadota* (mean 59 ± 13%), *Actinomycetota* (mean 14 ± 8%), *Bacteroidota* (average 13 ± 6%) and *Cyanobacteriota* (average 10 ± 8%) were dominant based on relative abundance (Supporting Information Fig. S12). *Pseudomonadota* (mean 17 ± 5%) and *Actinomycetota* (mean 7 ± 2%) were less abundant in the microplankton (3-90 µm) size fraction, instead *Cyanobacteria* (mean 29 ± 11%) and *Bacteroidota* (mean 23 ± 8%) were higher in relative abundance.

*Pelagibacterales* were high in relative abundance in the picoplankton across the estuary (mean 44 ± 13%), with lower relative abundances detected at NRE0 (mean 21 ± 11%, Fig. 6B, Supporting Information Fig. S13). Four genera of *Pelagibacterales* dominated the picoplankton, *Pelagibacter*, SYDM01 and IMCC9063 were abundant across all brackish stations, whereas *Fonsibacter* was predominantly present at NRE0. The dominant cyanobacterial genus was *Vulcanococcus* (*Synechococcus*-like) with 15 ± 9% and 8 ± 7% of relative abundance in the microplankton and picoplankton size fraction, respectively. On October 25 at NRE0, the class *Cyanobacteriia* contributed 31% of relative abundance to the microplankton and the potentially cyanotoxic cyanobacterial species *Planktothrix agardhii* accounted for 11.5% in relative abundance.

Principal Coordinate Analyses (PCoA) of prokaryotic and eukaryotic plankton community showed a clustering of NRE0 samples (Supporting Information Fig. S14). At station NRE0 33% and 18% of the ASVs of eukaryotic and prokaryotic plankton were unique, respectively. The prokaryotic plankton community at station NRE70 in July was more similar to the NRE0 communities sampled across summer and fall, in line with lower salinity and higher river discharge (Supporting Information Fig. S2, S13, S14). Across all samples, community composition clustered by size fraction rather than sampling time. The prokaryotic community at mid-estuary station NRE50 was usually distinct compared to up- and downriver stations.

### Connectivity between B-vitamins and plankton communities

Potential connections between B-vitamin/vitamer pools and the bacterioplankton and phytoplankton abundances were examined by correlation analysis (Fig. 4, Supporting Information Fig. S15). Dissolved OH-psB12 strongly and positively correlated with PC-SYN and PE-SYN abundance, additionally particulate OH-psB12 was positively correlated to PC-SYN. Similarly, dissolved B1 positively correlated with PC-SYN abundance (Fig. 4, Supporting Information Fig. S8B). Dissolved and particulate OH-psB12, and particulate B2 and B5 of the 0.2-3 µm size fraction positively correlated with bacterioplankton abundances. The (pico-)cyanobacterial order PCC-6307 was positively correlated with dissolved OH-psB12, particulate B1 and total psB12 of the picoplankton size fraction (Supporting Information Fig. S8, S15). Relative abundances of of picocyanobacteria (0.2-3 µm) were additionally positively correlated to DMB. The *Chlorophyta* genus *Bathycoccus* (0.2-3 µm) was positively correlated to dissolved OH-psB12 and total particulate psB12. In the 3-90 µm particulate size fraction the relative abundance of the Dinoflagellate genus *Levanderina* was positively correlated to total B12. In both size fractions *Polykrikos* was positively correlated to dissolved B1 and B3.

Redundancy analysis was used to examine if B-vitamin and vitamer concentrations significantly (p < 0.05) helped explain observed variability in plankton community composition. Select B-vitamins and vitamers together explained 30.6% of variation in eukaryotic plankton composition and 42% of the observed variance in prokaryotic plankton (Supporting Information Table S3). Specifically, particulate B3 and FAMP, and dissolved B1 and OH-psB12 were highly significant (p ≤ 0.005) explanatory variables for plankton community composition. Additional significant explanatory variables for the eukaryotic plankton community were dissolved B6 and particulate OH-B12. The strongest explanatory variable for the prokaryotic community was dissolved B1 and particulate B3 for eukaryotic plankton. Overall, multiple B-vitamins, B1- and B12-vitamers showed a statistically significant effect on plankton composition.

Salinity was the main driving environmental variable for eukaryotic and prokaryotic community composition, explaining 6.9 and 8.5%, respectively (Supporting Information Table S3). Measured environmental variables (aside from B-vitamins/vitamers) explained 13.9% and 20.4% of the observed variability in eukaryotic and prokaryotic plankton community composition, respectively. In both eukaryotic and prokaryotic communities, B-vitamin/vitamer concentrations cumulatively explained a greater proportion of variance, compared to hydrological and nutrient measurements during our summer–fall sampling.

## DISCUSSION

### Concentration patterns of B-vitamins and vitamers

Here, we uncover elevated concentrations of B-vitamins mid-estuary (Chl *a* max region) along with unique dissolved B-vitamin profiles associated with higher phytoplankton biomass. Key bacterio- and phytoplankton groups observed and their potential role as B-vitamin producers or consumers are disscued below and summarized in a conceptual figure in relation to the dissolved B-vitamins observed across the freshwater to polyhaline gradient of the NRE in summer and fall 2021 (Fig. 7). Our study identifies B1, B3, and B12 compounds as key compounds influencing prokaryotic and eukaryotic plankton communities. The simultaneous quantification of the two dissolved B12 forms, OH-B12 and OH-psB12, show that on average OH-psB12 (mean 0.9 ± 0.3 pM) was at about 47% of the concentration of OH-B12 (mean 1.9 ± 0.6 pM; Fig. 2A), supporting the hypothesis that psB12 could be a crucial source for B12-remodellers, especially in B12 limited systems. Dissolved OH-psB12 emerged as a key driver of both prokaryotic and eukaryotic community composition in the NRE, strengthening its importance as an indirect B12 source even for organisms thought to be unable to remodel the compound. Estimates suggest up to 17% of bacteria can salvage B12 (Shelton et al. 2019), and the genetic capability to remodel psB12 or use other cobalamin intermediates might be of high importance for planktonic microorganisms in the NRE and beyond. While OH-B12 is expected to be the dominant form of B12 in the sun-lit aquatic environments (Bannon et al. 2024a), OH-psB12 synthesized by cyanobacteria, could be an additional important source of B12 to organisms capable of remodeling psB12 (Helliwell et al. 2016). Since the 1950s/1960s evidence has accumulated highlighting B12 as an essential metabolite for aquatic microorganisms, however fewer than 40% of prokaryotes are predicted to synthesize B12 *de novo* (Shelton et al. 2019) and only now we are beginning to elucidate microbial sources, cycling, and transformations of the distinct B12 forms (Soto et al. 2023; Bannon et al. 2024a; Wienhausen et al. 2024).

**Fig. 7.**
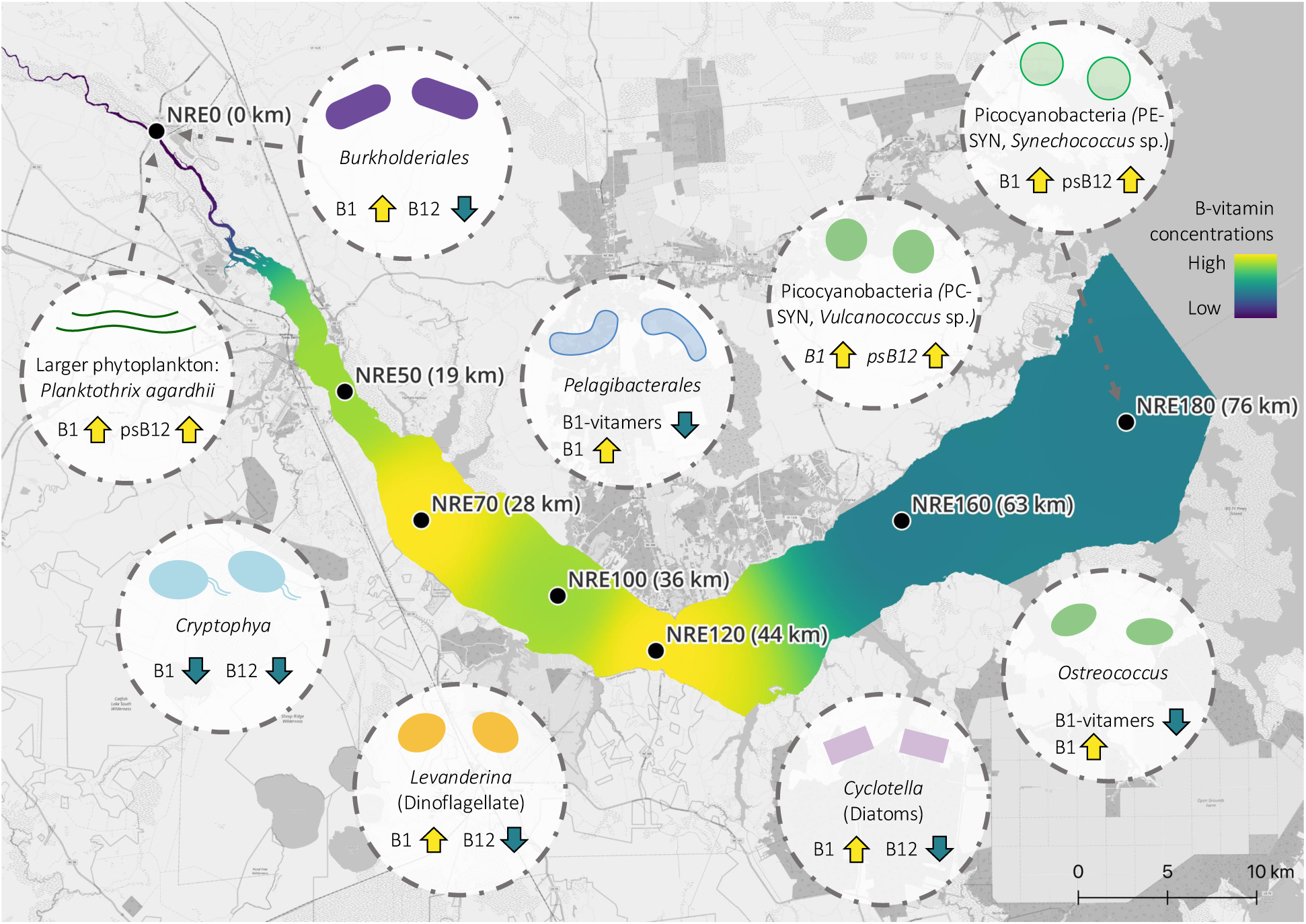
Conceptual map of key bacterio- and phytoplankton groups observed and their potential role as B-vitamin producers or consumers across the freshwater to polyhaline gradient of the NRE in summer and fall 2021. The color gradient of the estuary indicates the relative total dissolved concentrations of measured B-vitamins and vitamers. Yellow or blue arrows in the taxa bubbles indicate potential production (particulate & dissolved) or consumption (dissolved) of B-vitamins/vitamers, respectively, based on findings from culture studies and/or the correlation analyses presented and discussed here.

Tracking of B-vitamins/vitamer dynamics across the particulate and dissolved phase of an estuary shows that some of the compounds produced within an estuary are enriched in the dissolved phase compared to the particulate phase (FAMP, HET, OH-psB12; Fig. 5A). We hypothesize that these degradation compounds (FAMP, HET) and OH-psB12 are not readily used by most bacterio- and phytoplankton groups, due to a lack of sufficiently sensitive transporter or vitamin salvage pathways, and therefore accumulate. These compounds require further salvaging and remodeling activity to fulfil the cellular vitamin requirement and could provide an advantage for microorganisms with these metabolic pathways (Paerl et al. 2023a). For further discussion on B1, B3 and more details on the partitioning of B-vitamins and vitamers across the pico- and nano-/microplankton phases and the dissolved phase see the Supporting Information.

### Putative eukaryotic microbial sources and transformations of B-vitamins

Our data highlights a tight connection between dominant phytoplankton and specific vitamin profiles in the particulate pool, while we are still working to understand the vitamin availability resulting from algal blooms (and more broadly the exometabolomes of blooms over time and space). The phytoplankton growth in the middle of the estuary includes the biosynthesis of a phytoplankton-specific suite of vitamin/vitamers, and these micronutrients in turn can support growth of auxotrophic phytoplankton, including potential HABs, and bacterioplankton.

The dominant eukaryotic phytoplankton groups identified (with 18S rRNA gene sequencing) and their dynamics are typical for late summer, early fall in the NRE, consisting of diatoms, dinoflagellates, cryptomonas, and chlorophyta (Gong and Marchetti 2019; Gong et al. 2020). The highly abundant *Cyclotella* (diatom) could have a B12 requirement and is likely prototrophic for B1, based on findings of experiments with one isolate (Tang et al. 2010; Bertrand and Allen 2012), however B12 auxotrophy has been shown to be strain specific. Contrastingly, *Cryptophytes* can have a requirement for both exogenous B1 and B12 (Tang et al. 2010), making them likely consumers of B1, B12, and their respective vitamers. Thus, surges of *Cryptophyta* (Cryptophytes) in NRE could be governed by the availability of B1 and B12. Similarly, the abundance of *Chlorophyta* taxa *Ostreococcus*, *Micromonas*, *Bathycoccus* could be regulated by the availability of B1 vitamers, as they are B1 auxotrophs but can salvage B1 from vitamers (Paerl et al. 2015, 2023b). Thereby, these taxa simultaneously consume vitamers and potentially function as a source for ‘regenerated’ vitamin in the upper estuary (NRE50, 70; 25 Oct).

*Levanderina* (Dinoflagellate) recurrently surged during summer/fall (Supporting Information Fig. S9) and, based on elevated microplankton particulate B1 concentrations during higher abundances of *Levanderina* (e.g. 28.1 pM B1 and 21.8% relative abundance of *Levanderina fissa* NRE70 25 Oct), these dinoflagellates are potential B1 producers and sources of B1 to higher trophic levels. Previously, B1 biosynthesis transcripts (*thiC*, *thiE*) were abundant during a dinoflagellate bloom (*Levanderina fissa*) in the NRE (Gong et al. 2017), further supporting that some bloom-forming Dinoflagellates in the NRE are significant B1 producers.

### Insight into B-vitamin cycling along an estuary continuum

Our measurements have revealed several new perspectives on vitamin cycling within (temperate, long residence time) estuaries: (1) freshwater input indirectly fuels increased levels of vitamins mid-estuary by promoting algal growth and (2) B-vitamins (dissolved and particulate) are likely utilized quickly in the lower estuary and transferred to higher tropic levels in the benthos or downstream (Fig. 7), similar to macronutrients. During extended periods of high discharge, the mid estuary peak of planktonic biomass (Chl *a,* bacterial abundance, POC) could shift further downstream or not occur and accordingly affect vitamin supply to the larger Pamlico Sound (Paerl et al. 2010; Hall et al. 2013). Surprisingly, B-vitamin/vitamer concentrations were not overtly elevated for all compounds in the freshest region of the NRE as hog and poultry operations are extensive within the Neuse River watershed (Lebo et al. 2012) and both are established sources of nutrients and presumably B-vitamins (Lunetta et al. 2022).

While we find evidence that estuaries can be a source of B-vitamins/vitamers to downstream coastal waters (Fig. 2, 4), the extent of supply is likely dependent on the freshwater discharge and vitamin-specific (Fig. 7), as some B-vitamins or vitamers appear to accumulate in the dissolved phase (e.g. psB12, FAMP, Fig. 5A). These accumulated compounds could function as more ‘recalcitrant’ vitamin/vitamers for the phytoplankton and bacterioplankton downstream. Overall, the oligohaline water of the NRE was lower in B-vitamins and characterized by distinct plankton and B-vitamin/vitamer profiles compared to the brackish part of the estuary (see Supporting Information). Prior work found inverse correlations between B-vitamin concentration and salinity suggesting rivers and groundwater as sources of B-vitamins to coastal systems (Gobler et al. 2007). Data from coastal regions influenced by the Amazon (Brazil; dissolved B1 and B6) and Moulouya (Morocco; dissolved B1, B2, B6, B12) rivers found that estuaries were not a major source to the coastal ocean (Barada et al. 2013; Tovar-Sánchez et al. 2016). While we find negative correlations of dissolved B2 and B6 with salinity, positive correlations are evident with dissolved compounds (HMP, B5 and OH-psB12), indicating that a general pattern for dissolved B-vitamins with salinity might not exist in this system. Importantly, salinity is a driving factor for bacterio- and phytoplankton community structure in the NRE, as shown here and in previous studies (Paerl et al. 2020; Sánchez-Gallego et al. 2025), thereby salinity indirectly affects B-vitamin concentrations. Overall, our data indicate that the NRE delivers B-vitamins and vitamers into Pamlico Sound (massive component of the APES) but the extent is expected to depend on flushing (antecedent precipitation) and changes in microbial community composition. Moreover, growth and proliferation of HAB spp. within the Pamlico Sound may be impacted by B-vitamin/vitamer delivery via the NRE – making the process impactful to food web, water quality, and ecosystem health.

### Temporal dynamics of B-vitamins and vitamers

B vitamin concentrations across the estuary were highly dynamic - from sub picomolar to high picomolar levels - revealing strong short-term variability driven by synthesis, uptake, and degradation processes, as well as sporadic surges in pico- and microplankton populations. Two modes of B-vitamin dynamics were observed: 1) moderate concentration dynamics (2 to 4-fold change) within a compound specific range (e.g. dissolved HMP; particulate FAMP) including occasionally elevated concentrations and 2) strong changes in concentrations (10-fold or more) characterized by occasionally high peaks in concentration (e.g. dissolved B5 and cHET; particulate B2; Fig. 3, 4). We argue that the dynamic patterns observed with mode 1 reflect oscillations between states of balance/imbalance for the planktonic communities due to vitamin/vitamer production and consumption, resulting in slight to moderate fluctuations. We hypothesize that mode 2 is driven by the strong rise and fall (or elevated activity) of specific planktonic populations leading to distinct vitamin profiles (e.g. October 25^th^ NRE0, *Planktothrix agardhii*, peak particulate psB12).

During a bloom, particulate B-vitamin and vitamer concentrations can increase sharply, potentially disproportionally to changes in the dissolved pool as observed for (microplankton) particulate B-vitamin and vitamer concentrations at NRE0 on 25 Oct. These sporadic episodes of rapid biomass accumulation likely reflect phases, during which intracellular vitamin synthesis (or uptake and salvage) dominates over release. As phytoplankton become nutrient-limited or physiologically stressed, however, extracellular release (and leakage) of vitamins can rise, a pattern documented for DOM in both cultures and environmental samples. While the transfer into the dissolved phase might be taxa specific (Sultana et al. 2023), the observed vitamin concentration patterns suggest overall minimal transfer of vitamin/vitamer to the communal dissolved pool – outside of declines and death.

### B-vitamins/vitamers shape microbial community composition

Here, we find evidence of B-vitamins - especially dissolved B1 and OH-psB12 and particulate B3 and FAMP - as key explanatory variables for both prokaryotic and eukaryotic plankton estuarine communities (Supporting Information Table S3). This is congruent with widespread auxotrophy for B1 and B12 and the important role of B3 in cellular metabolism. Together, this builds a greater perspective that these compounds are key ‘shapers’ of plankton communities. The concentrations of B-vitamins and vitamers collectively explained 42% and 31% of the observed variance in the prokaryotic and eukaryotic planktonic community respectively - highlighting the overall importance of B-vitamins in shaping microbial community composition in estuarine waters, complementary to previous observations from bottle/mesocosm incubations that have demonstrated vitamin-driven changes in plankton communities in certain habitats (reviewed in: Bertrand & Allen 2012; Joglar et al. 2020).

In the NRE, Picocyanobacteria are likely *de novo* synthesizers of B1 and psB12 and reach higher abundances in brackish waters away from the freshwater endmember of the NRE (e.g. NRE0; Paerl et al. 2020). The detection of peak particulate Me-psB12 (3.4 ± 0.6 pM) in the microplankton on 25 October coincides with a high relative abundance of the filamentous cyanobacteria *Planktothrix agardhii* in the microplankton size fraction. Congruently, we found a link between dissolved B1, OH-psB12 concentrations in the picoplankton, and dissolved OH-psB12 with the abundance of PC-SYN (Fig. 4, Supporting Information Fig. S8, S15) highlighting them as sources. As dissolved OH-psB12 is not readily available to most plankton a tighter coupling between picocyanobacterial abundance and psB12 is observed compared to dissolved B1, which is likely rapidly utilized. Recent field data (Roskilde Fjord Denmark; Northwest Atlantic) also point to picocyanobacteria as important B1 and psB12 sources (Bannon et al. 2024b; Bittner et al. 2024). These lines of evidence support that picocyanobacterial abundance could function as a proxy for psB12 concentrations, and that psB12 provided by (pico-) cyanobacteria could significantly promote taxa capable of cobalamin remodeling (Helliwell et al. 2016; Soto et al. 2023). Salvage of B12 from degraded cobalamin and DMB may represent an important yet overlooked pathway in aquatic systems, as cyanobacteria are already considered key planktonic community members - especially through C and N-fixation - their role may extend further as producers of key vitamins such as B1 and psB12.

## Author contribution statement

**MJB:** Conceptualization, Methodology, Formal analysis, Investigation, Data Curation, Writing – Original Draft, Writing – Review & Editing, Visualization, Funding Acquisition

**CCB:** Methodology, Investigation, Data Curation, Writing – Review & Editing

**ER:** Methodology, Investigation, Data Curation, Writing – Review & Editing

**GL:** Investigation, Writing – Review & Editing, Visualization

**EMB:** Methodology, Resources, Writing – Review & Editing, Funding Acquisition

**LR:** Writing – Review & Editing, Supervision, Funding Acquisition

**RWP:** Conceptualization, Methodology, Investigation, Resources, Data Curation, Writing – Review & Editing, Supervision, Funding Acquisition

## ACKNOWLEDGEMENTS

This work was supported by Independent Research Fund Denmark (9040-00067B to LR, RWP, and Anders F. Andersson). MJB received funding from the European Union’s Horizon 2020 research and innovation programme under the Marie Skłodowska-Curie grant agreement No 801199. RWP acknowledges support from NSF OCE NSF OCE awards Oand 2416286. EMB acknowledges support from NSERC Discovery Grant RGPIN-2015-05009 and Simons Foundation Grants 504183 and 1001702. We are grateful to Jeremy Braddy and Amy Bartenfelder for assistance with water collections. We thank the whole UNC-IMS MODMON team for analyzing and providing hydrological, chemical, and biological data. We thank Malcolm Barnard for help with filtrations. We thank UNC-IMS for providing lab space and local logistical support. We thank Anders F. Andersson for input on the study.

## CONFLICTS OF INTEREST

No conflicts of interest.

## DATA AVAILABILITY STATEMENT

The data that support the findings of this study are openly available in the Supporting Information Data S1, S2 and S3 deposited at https://doi.org/10.11583/DTU.31353040. Raw sequence reads from 16S and 18S rRNA genes are deposited at NCBI under accession PRJNA1175993.

## Notes

### Competing Interest Statement

The authors have declared no competing interest.

