## Supplementary Information for "Novel connections between B-vitamins and microbial communities along biogeochemical gradients in a large temperate estuary"

**Contents of this file:**

Supporting Methods

Supplementary Text

Supporting Figures 1 to 10

Supporting Tables 1 to 3

Caption for Supporting Information Data S1 to S3

References for Supporting Information

**Supporting Data for this manuscript:** https://doi.org/10.11583/DTU.31353040

Data S1

Data S2

Data S3

**Supporting Methods - Metabolite analysis**

To measure the extracted metabolites on the Dionex Ultimate-3000 LC system coupled to a TSQ Quantiva triple-stage quadrupole mass spectrometer (ThermoFisher) samples were resuspended in HPLC buffer A (20 mM ammonium formate, 0.1 % formic acid), diluted (for dilution factors see Supporting Information Table S2) and a volume of 5 µl was injected. Quality Control (QC) pools were created for each matrix group by pooling equal portions of each sample from the group. Metabolites were quantified by using standard addition method. Standards were run for each group separately in the QC-pool, resulting in specific limits of detection (LOD) and limits of quantification (LOQ) per group (Supporting Information Data S2). Measurements with concentrations between LOD and LOQ were manually inspected and analyzed in a batch per batch method (Boysen et al. 2018).

Due to issues upon injection to the mass spectrometer, data of only one injection was available for two particulate samples and included in the analysis (NRE180 replicate A 26 Jul, NRE70 replicate A 13 Sep). Three measurements of particulate B1 and B2 were excluded as concentrations were more than 10 x higher than other respective biological replicates (0.2-3.0 µm NRE70 13 and 27 Sep, 3.0-90 µm NRE50 27 Sep). Measurements of dissolved metabolites from 25 Oct NRE70 were based on two biological replicates as one replicate had to be removed due to odd chromatography. Additionally, six samples were lost during processing, which resulted in no dissolved metabolite measurements for NRE70, and only one replicate for NRE100 and only two NRE0 replicates from 11 Oct.

**Supplementary Text**

**Distinct B-vitamin patterns in oligohaline estuarine waters**

Generally, freshwater station NRE0 exhibited lower phytoplankton biomass (POC, *Chl a*) and bacterioplankton abundances, but higher dissolved inorganic nitrogen and silicate concentrations and elevated turbidity. Except on 25 October when concentrations of POC, *Chl a* and primary production peaked at NRE0. Overall, station NRE0 was characterized by lower B-vitamin and vitamer concentrations compared to brackish stations, except maximum concentrations of particulate B-vitamins (B5, psB12 and HET) in the microplankton on 25 October (Fig. 3B). Distinct B-vitamin/vitamer profiles were associated with NRE0 during non-bloom and bloom samples, contrasting to brackish stations, likely a result of distinct environmental characteristics and the freshwater specific picoplankton and microplankton community, lower B1 and B12 production was previously observed in select freshwater phytoplankton cultures compared to marine phytoplankton cultures (Carlucci & Bowes 1970, Nishijima et al. 1979).

**Unique putative B-vitamin sources in freshwater**

Pico- and microplankton communities were distinct at the freshwater station (NRE0) compared to the brackish stations and included freshwater-specific taxa e.g. the widespread freshwater genus *Fonsibacter (Pelagibacterales* affiliate) (Tsementzi et al. 2019). Additionally, higher relative abundances of heterotrophic *Burkholderiales* were characteristic at NRE0, implicated as core B1 synthesizers in a Danish fjord (Bittner et al. 2024), highlighting *Burkholderiales* as potential *de novo* B1 synthesizers at NRE0, as *Pelagibacterales* are pyrimidine auxotrophs. Elevated inorganic nutrient concentrations at NRE0, could promote larger phytoplankton populations (Paerl et al. 2020) that could function as *de novo* B1 synthesizers at NRE0, supported by the spatial distribution observed for particulate B1, with a higher proportion of B1 within the microplankton populations in the upper estuary compared to the picoplankton. This suggests surges in abundance of prototrophic microplankton could be an important intermittent source of B1 in freshwater systems.

Freshwater samples from 25$^{th}$ of October showed peak primary production and *Chl a*, coinciding with a surge in relative abundance of the non-diazotrophic and potentially toxic *Planktothrix agardhii* (Chaffin et al. 2018). Such an event is uncommon at NRE0, typically phytoplankton blooms occur in the middle of the estuary (Pinckney et al. 1997).

Simultaneously we detected the fungal order *Rhizophydiales*, which is an obligate parasite for *Planktothrix agardhii* (McKindles et al. 2021). Parasitic fungi indirectly affect the availability of organic matter in aquatic systems (Grossart et al. 2019); possibly parasitic fungi also alter vitamin availability (via cell death) and impacted the specific vitamin profiles observed on 25$^{th}$ of October (peak concentrations of B1, B3, B5 and OH-psB12 in the microplankton biomass). Our data indicates that prototrophic filamentous cyanobacteria could be important B-vitamin sources (to dissolved and particulate phases) for other microorganisms and higher trophic levels in aquatic environments as picocyanobacteria (PE-SYN, PC-SYN) were present in lower abundances at NER0 compared to the brackish stations.

**Concentrations patterns of B-vitamins and vitamers**

Our discussion will focus on the dynamics of B1, B3 and B12 compounds as these were identified as key compounds for both prokaryotic and eukaryotic plankton communities. Dissolved B1 concentrations (19 to 74 pM) in NRE indicate short-term (weeks) temporal variation, as previous measurements of dissolved B1 from NRE0 and NRE180 (11 Nov 2021) were around 70 pM (Paerl et al. 2023a). Our dissolved B1 measurements add to similar measurements from a Danish brackish system and the North Sea (Bruns et al. 2023; Bittner et al. 2024). Contrastingly, particulate B1 concentrations were about 10-fold higher for each size fraction than previously reported from Denmark, in line with up to 10-fold higher POC in NRE. When normalized to POC, particulate B1 was in a similar range suggesting a possible general ratio of around 210 pmol particulate B1 to 1 mol of POC in coastal brackish systems (NRE: 2.9 ± 2.1 x 10-$^{7}$ mol B1 per mol POC; Roskilde fjord: 1.3 ± 1.0 x 10-$^{7}$ mol B1 per mol POC (Bittner et al. 2024). The detection of B1 degradation compounds FAMP in both picoplankton and microplankton, further supports that intracellular B1 degradation or FAMP uptake occurs in both planktonic size fractions (Paerl et al. 2023a). Use of B1-vitamers could help stabilize B1 stocks within cell, similar to recent findings, we find signs of stronger dynamics for dissolved B1 vitamers (AmMP, HMP, cHET and HET) than for B1 (Bittner et al. 2024, Longnecker et al. 2024).

B3 is the essential precursor for NAD/NADH or NADP/NADPH and therefore required for the synthesis of other B-vitamins (Kirkland & Meyer-Ficca 2018) but its distribution and role in planktonic microbial communities remains mostly unknown. Here, we measure one form of B3, niacinamide, therefore probably underestimating the concentration of total bioavailable B3 in the particulate and especially the dissolved phase, as niacin is the expected dominant form (Aguilera-Méndez et al. 2012). The NRE exhibited a wide concentration range for dissolved niacinamide (45-513 pM), if similar quantities of niacin are present and on what scales these two forms of B3 are transformed in the environment remains unknown. Limited dissolved B3 measurements indicated niacin concentrations below 50 pM in winter and spring at a coastal site of the North Sea (Bruns et al. 2022, 2023), whereas niacin concentrations across seasons in the Northwest Atlantic showed a broad concentration range of niacin from 11 to 319 pM (Bannon et al. 2025). Particulate niacinamide measurements from the Northwest Atlantic were on a concentration range of 3 to 124 pM (Bannon et al. 2025), size-fractionation of the particulate phase now allows us to show that the broad concentration range of niacinamide found in NRE was predominantly found in the picoplankton (14 to 306 pM), highlighting that picoplankton populations might be dominant producers and consumers of B3, providing a first glimpse into the unexplored aquatic B3 cycle.

The range of dissolved OH-B12 concentrations in the NRE is similar to concentration ranges reported from the Mediterranean Sea (0.09-3.13 pM, (Suffridge et al. 2018) and the Atlantic open ocean (NW: 0.10-2.50 pM (Suffridge et al. 2017, Bannon et al. 2025); Eastern: 3.1±0.2 pM). The simultaneous quantification of five cobalamin forms in the particulate phase reveals unforeseen partitioning in particulate phases. Typically, Ado-B12 was the dominant biologically active form of B12 in the picoplankton, which presumably includes most prokaryotic organisms capable of B12 *de novo* synthesis or salvage. Both Ado-B12 and Me-B12 were mostly below the detection limit in the microplankton as most eukaryotic phytoplankton require an exogenous source of B12 and would be likely to take up OH-B12 from the sun-lit ambient water. The detection of particulate DMB in both the pico- and microplankton provides evidence of potential DMB production or accumulation that would support B12 salvaging or remodeling activity. In both the pico- and microplankton particulate phases, DMB concentrations were more than 10-fold higher in the NRE than the only reported measurements of environmental particulate DMB from the Northwest Atlantic (0.13 ± 0.11 pM; 0.01-0.52 pM)(Bannon et al. 2025), but with similar DMB/POC values, as POC concentrations were about 10-fold lower in the Northwest Atlantic.

**B-vitamins across dissolved and particulate phases**

Concentrations of dissolved and particulate B-vitamins/vitamers that fall closely around the identity line (1:1 ratio) are indicative of equal concentrations in the dissolved and particulate phase. This was generally the case for ratios of particulate to dissolved B1, B3 and

B5 (Fig. 5A). While some dynamics in the phase partitioning can be observed, indicating short-term temporal variation in the balance of supply and demand, the overall ratios close to the identity line imply a balance between for example dissolved B1 and B1 required by the planktonic biomass. Positive correlations between dissolved B1, HMP, and HET suggest tight cycling of these compounds through synthesis, degradation, release, and/or B1 salvage. Other B-vitamins and vitamers (B2, FAMP, HET and OH-psB12) showed higher concentrations in the dissolved phase than in the particulate phase, indicating an excess in availability. Presumably, degradation compounds (FAMP and HET) can be abiotically formed in the dissolved phase for example via photodegradation, oxidation or temperature (Lukienko et al. 2000, Hanson et al. 2016). Contrastingly, in most samples OH-B12 concentrations were higher in the particulate phase than the dissolved phase, indicating a possible imbalance between B12 supply/demand and rapid assimilation of dissolved OH-B12.

**B-vitamins across pico- and nano-/microplankton phase**

While some dynamics in the partitioning between the two particulate size fractions can be observed, the overall ratios of multiple vitamins (B1, OH-B12, B2, B5) group close to the identity line, highlighting that the stock of vitamins in each size fraction is similar and vitamins do not accumulate in the dissolved phased but are potentially available to higher trophic levels when bound in planktonic biomass. This suggests a tight coupling of *de novo* B12 synthesis in bacterioplankton and B12 uptake/acquisition by eukaryotic microplankton with a B12 requirement. B12 related compounds DMB and OH-psB12 were typically enriched in the picoplankton biomass compared to microplankton biomass, as picocyanobacteria are abundant in the NRE (Paerl et al. 2020) and likely the main source of psB12 (Fig. 3, Supporting Information Fig. S5). Some bacterioplankton and microplankton populations are potentially capable of exchanging the lower ligand of psB12 with DMB (Helliwell et al. 2016, Wienhausen et al. 2024). Overall, our simultaneous quantification of dissolved and size-fractionated particulate B-vitamins will be useful to further budget environmental pools of vitamins and their exchanges and to trace their flows in aquatic food webs.

**Connectivity of B-vitamins and vitamers**

Most vitamin or vitamer concentrations were not correlated across phases, indicating that abiotic and biotic transformations occur between phases (Supporting Information Fig. S7). One exception was the positive correlation of particulate B2 and DMB in the picoplankton biomass, B2 serves as a precursor for DMB synthesis via the BluB enzyme (Taga et al. 2007), thereby linking these two compounds. Occasionally picoplankton vitamin profiles clustered with the corresponding microplankton vitamin profile (NRE0 27 Sep and 11 Oct, NRE120 11 Oct, NRE70 25 Oct; Fig. 3B), indicating possibly a result of picoplankton cell aggregation or trophic transfer of vitamins. At other times particulate vitamin profiles of the same size fraction of adjacent stations grouped, indicating occurrence of planktonic populations across parts of the estuary and a downstream transport of planktonic biomass including vitamins (e.g. 0.2-3 µm NRE100 and NRE120 11 Oct; Fig. 3B).

**Supporting Figures**

A

B

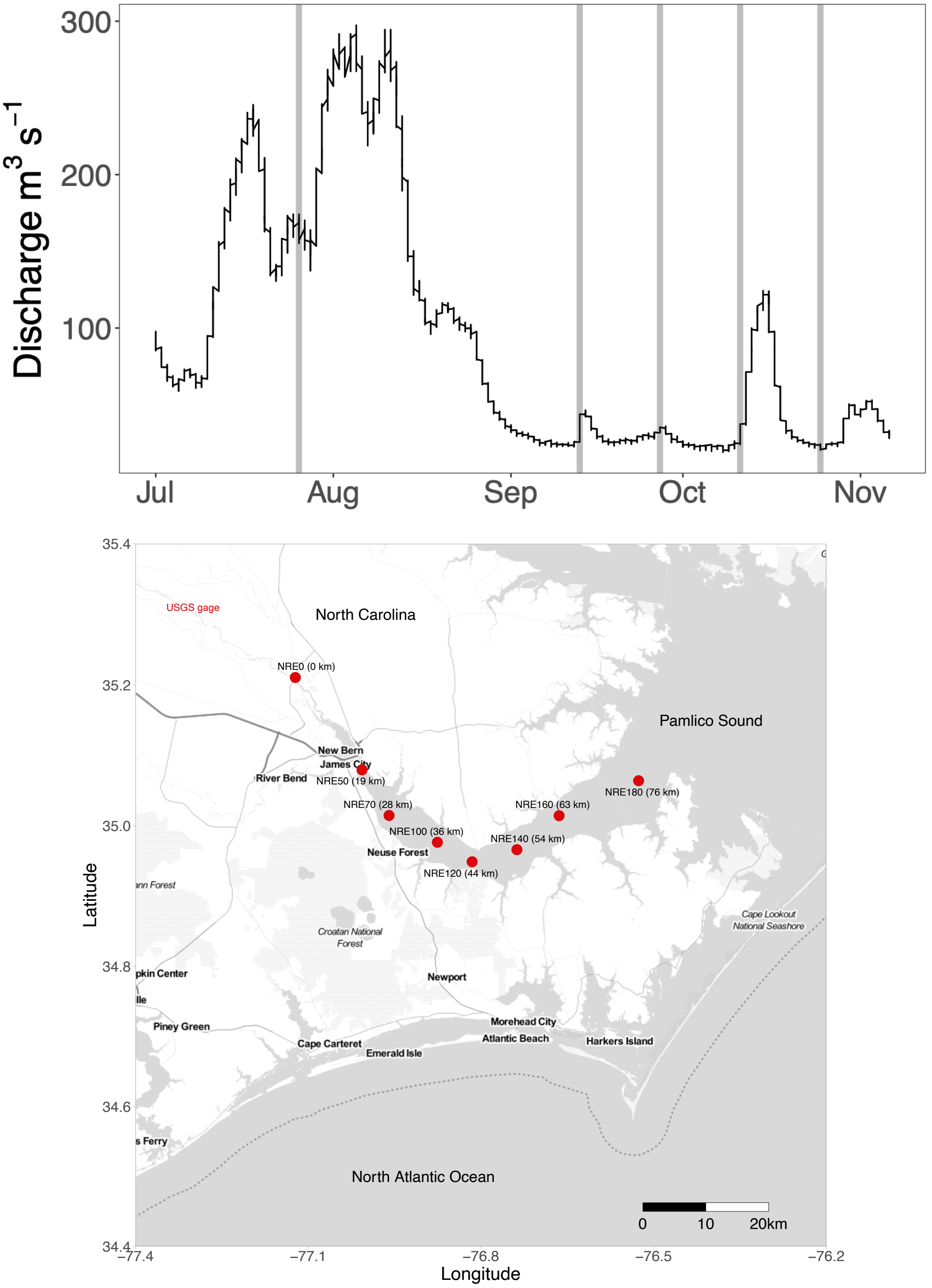


**Fig. S1.** Discharge measurements from USGS gauge 02091914 (**A**) in Neuse River near Fort Barnwell, NC. Vertical gray lines indicate days of sampling in NRE. Data was retrieved from the USGS National Water Information System Web Interface https://waterdata.usgs.gov/nwis. Location of USGS gauge indicated on map (**B**). NRE map was generated with ggmaps (Kahle & Wickham 2013) with the aid of packages ggsn and ggrepel.

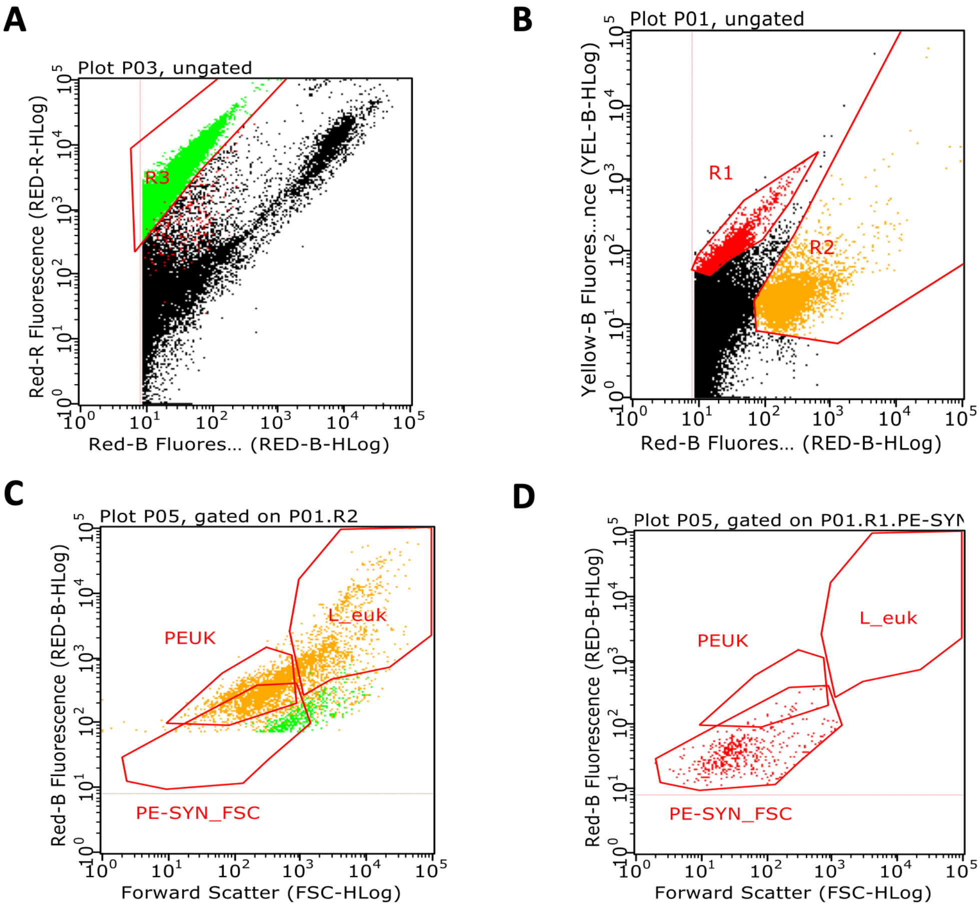


**Fig. S2.** Example flow cytograms and gating used in analysis of NRE samples to determine phytoplankton morphotypes. In (**A**) PC-SYN (gate R3) and other cells are distinguished based on red fluorescence from red excitation light (Red-R) versus red fluorescence from blue excitation light (Red-B), as PC-SYN cells exhibit a higher Red-R signal relative to Red-B. Initial gating of PE-SYN (R1) and eukaryotic was done based on yellow fluorescence from blue excitation (Yellow-B) versus Red-B (**B**), then (using hierarchical gating) picoeukaryotic phytoplankton (PEUK), larger eukaryotic phytoplankton (LEUK), and PE-SYN_FSC (single cells) were distinguished based on red fluorescence from blue excitation light (Red-B) and forward scatter (FSC) (**C**, **D**).

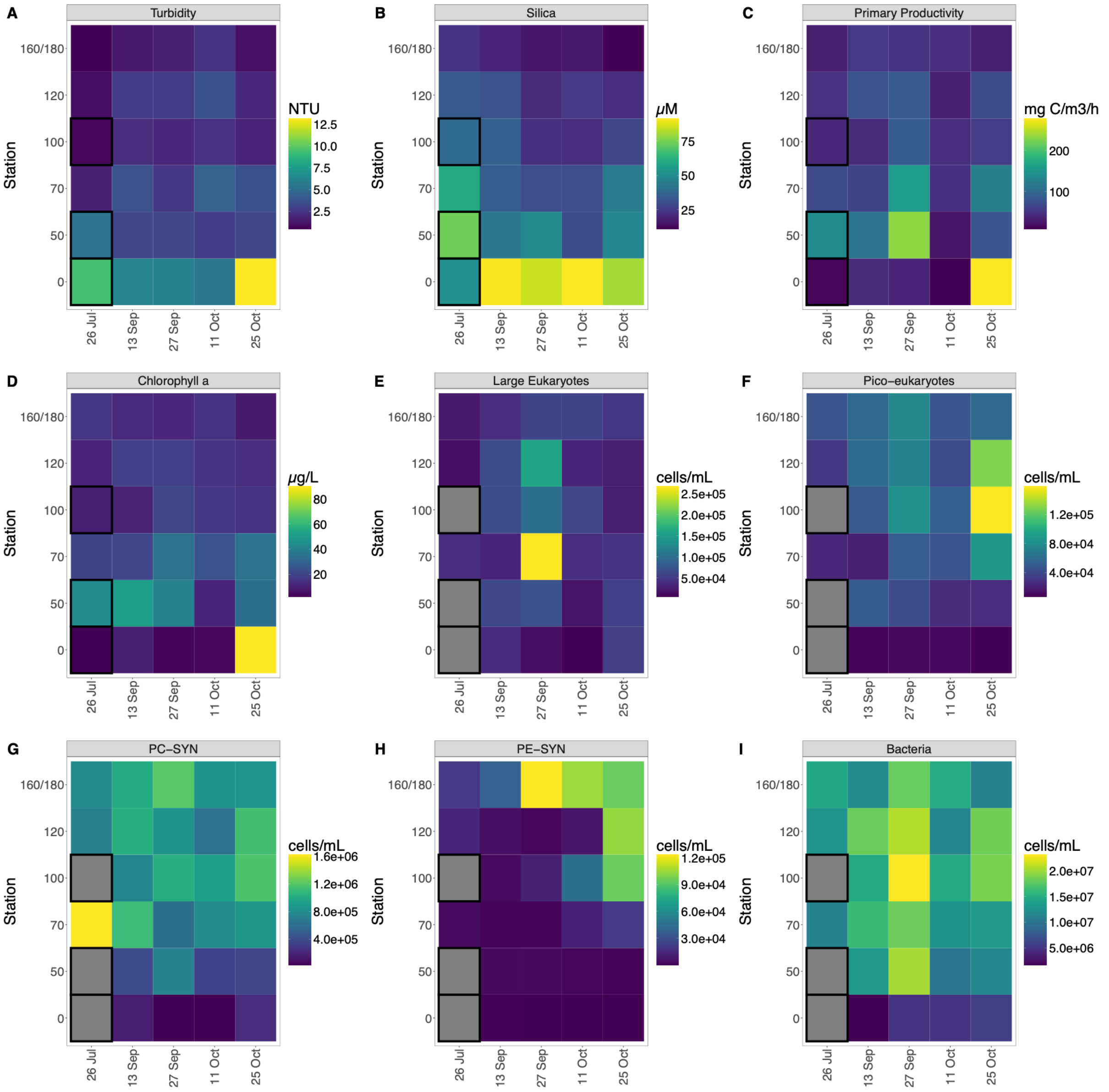


**Fig. S3.** Measurements of turbidity (**A**), Silica (**B**), Primary Productivity, Chlorophyll a (**D**), phytoplankton morphotypes (**E-H**) and bacterial cell abundance (**I**) determined by flow cytometry. PE-SYN: *Synechococcus*-like phycoerythrin-rich cells. PC-SYN*: Synechococcus*-like phycocyanin-rich cells. Black outlines indicate stations were not sampled for vitamin and plankton analysis.

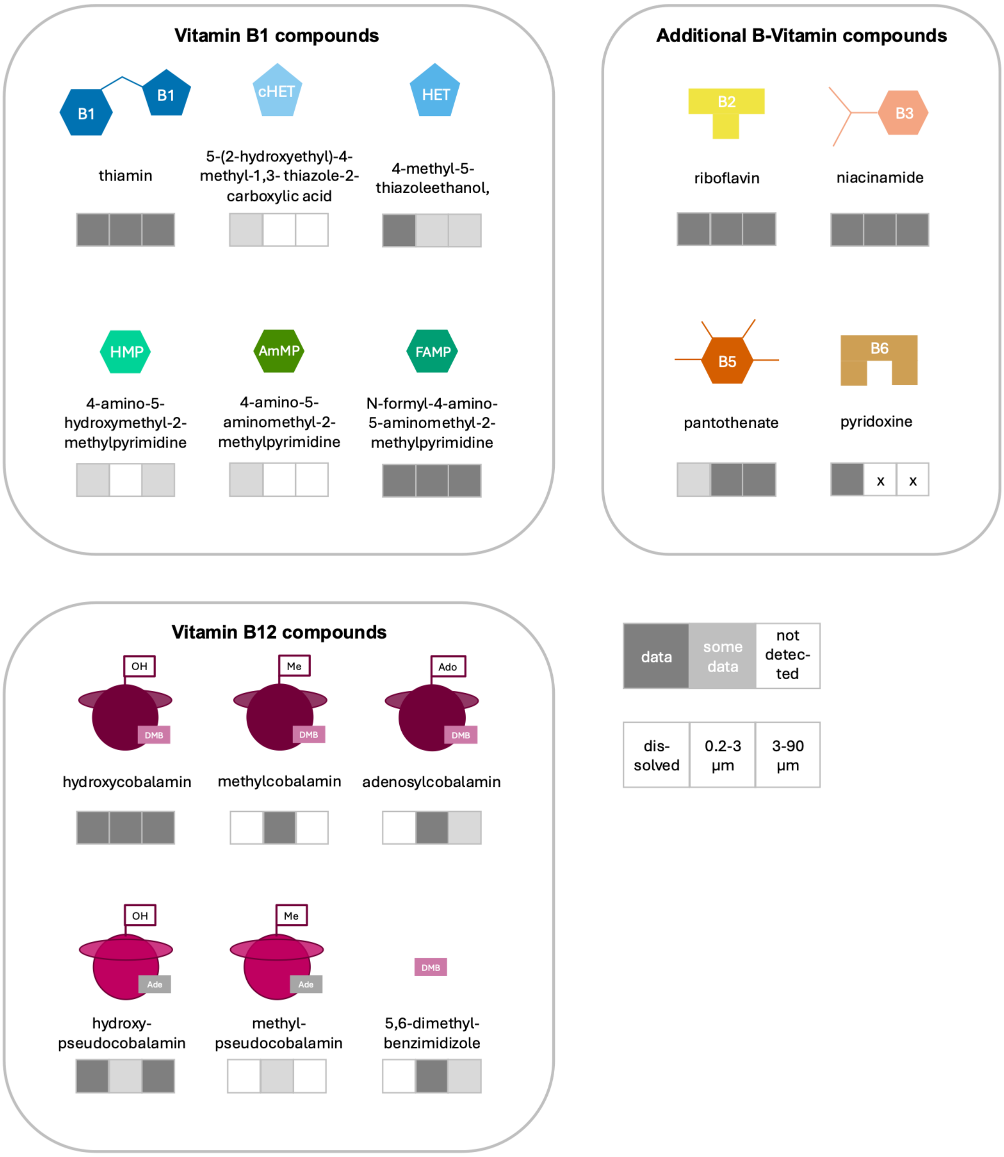


**Fig. S4.** Schematic overview of analytically targeted compounds and their available B-vitamin/vitamer measurements across the three metabolite datasets: dissolved (left box), pico-particulate (0.2-3 µm, middle box) and nano-/micro-particulate (3-90 µm, right box). The gray scales below schematic compounds indicate the extent of available measurements, if >5 samples were below LOD or LOQ, the classification 'some data' was used and if <5 measurements were available for a given compound in a certain dataset 'not detected' was used. For further details per matrix grouping see Supporting Information Data S2 and for compound measurements see Supporting Information Data S3. B6 was not targeted in the particulate size fractions, indicated with an x.

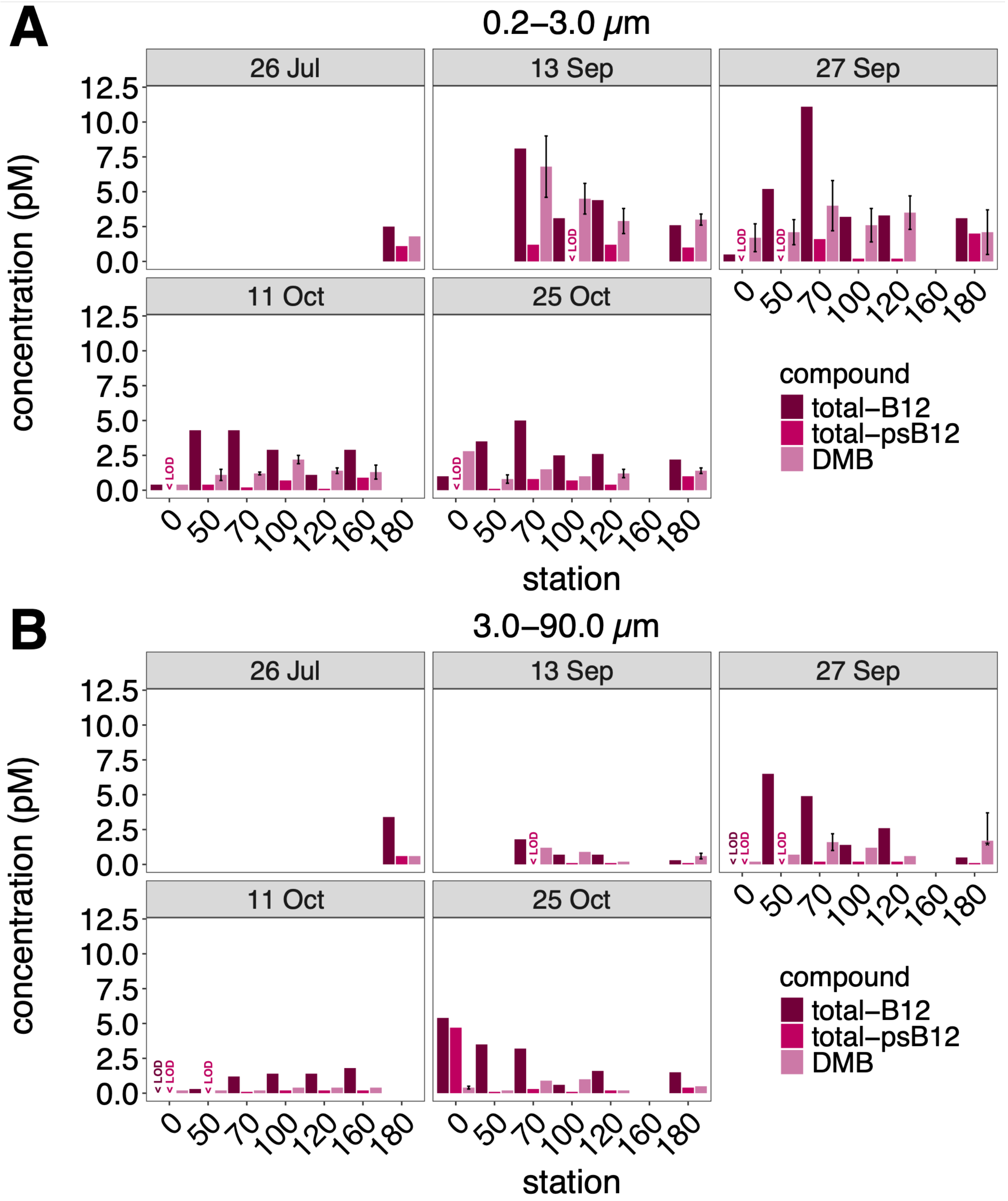


**Fig. S5.** Bar plot of particulate total B12, total-psB12 and DMB from the 0.2-3 μm (**A**) and 3.0-90.0 μm size fraction (**B**). DMB error bars show ± standard deviation of biological replicate water samples, on Sept. 27th at NRE180 the standard deviation is higher than the mean concentration across three replicates, indicated with an asterisk.

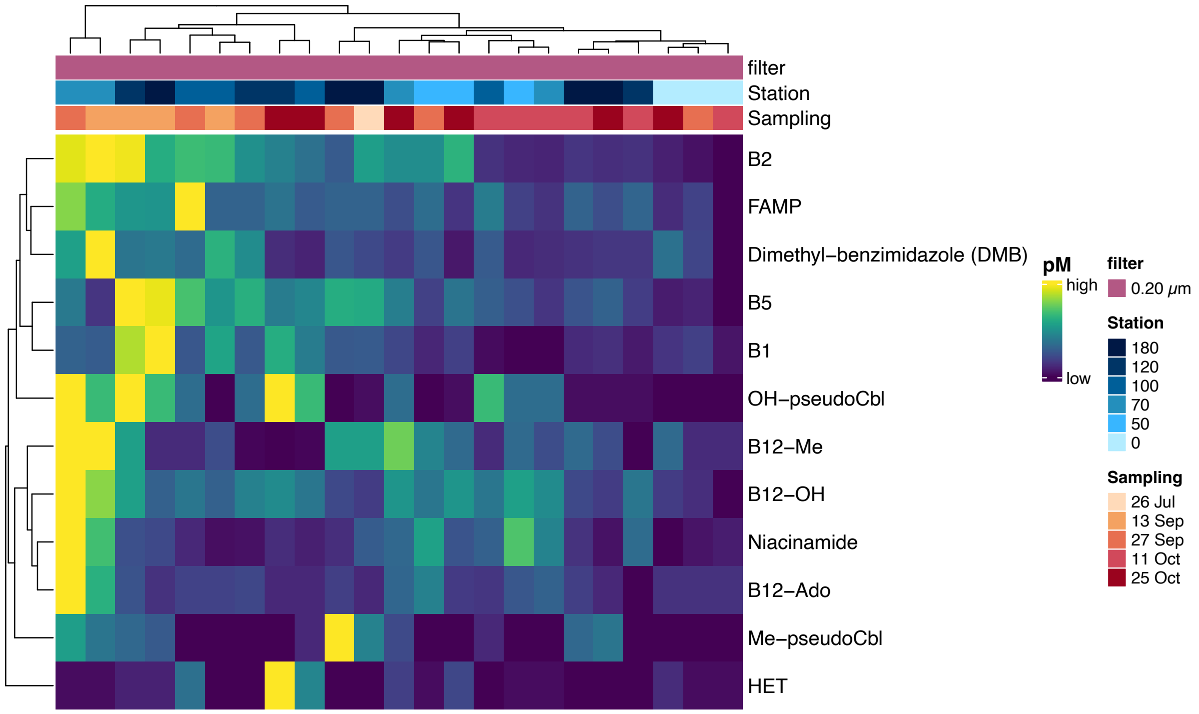


**Fig. S6.** Heat map of particulate B-vitamins and vitamer profiles from the 0.2-3 µm size fraction. Columns and rows were clustered based on Euclidean distances corresponding to differences between samples and relative metabolite concentrations, respectively. Metabolite concentrations are relative to the concentration range of each compound. The station and sampling time point are indicated on the top. For measurements below limit of detection or limit of quantification, the concentration limits are displayed (Supporting Information Data S2A).

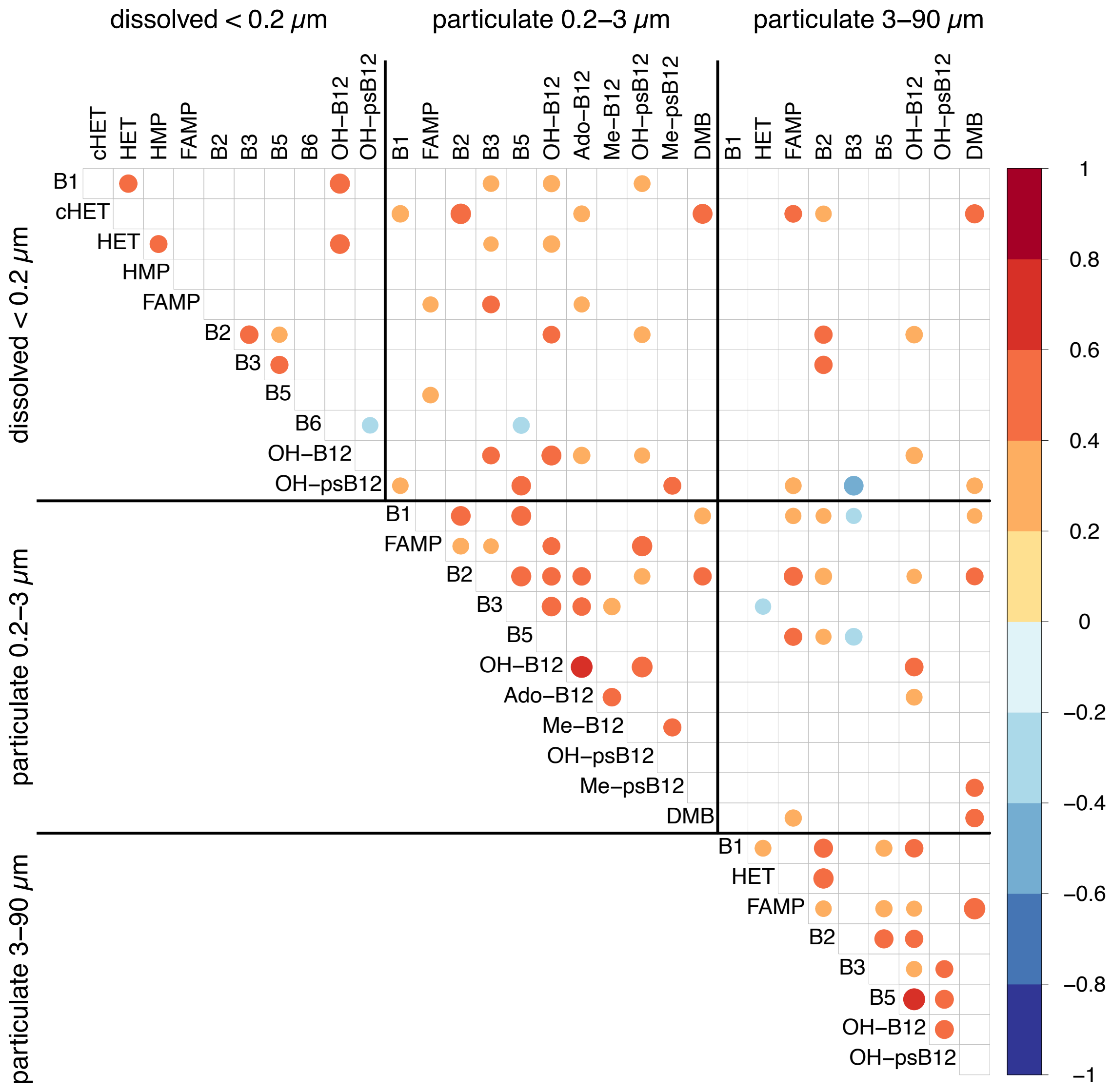


**Fig. S7.** Correlation plot of significant ($p < 0.05$) Kendall's rank correlations between metabolites from dissolved and particulate samples. For abbreviation of metabolite names see Supporting Information Fig. S4 or Data S2. Metabolites below of limits of quantification or detection were replaced by the lower LOD prior to correlation analysis.

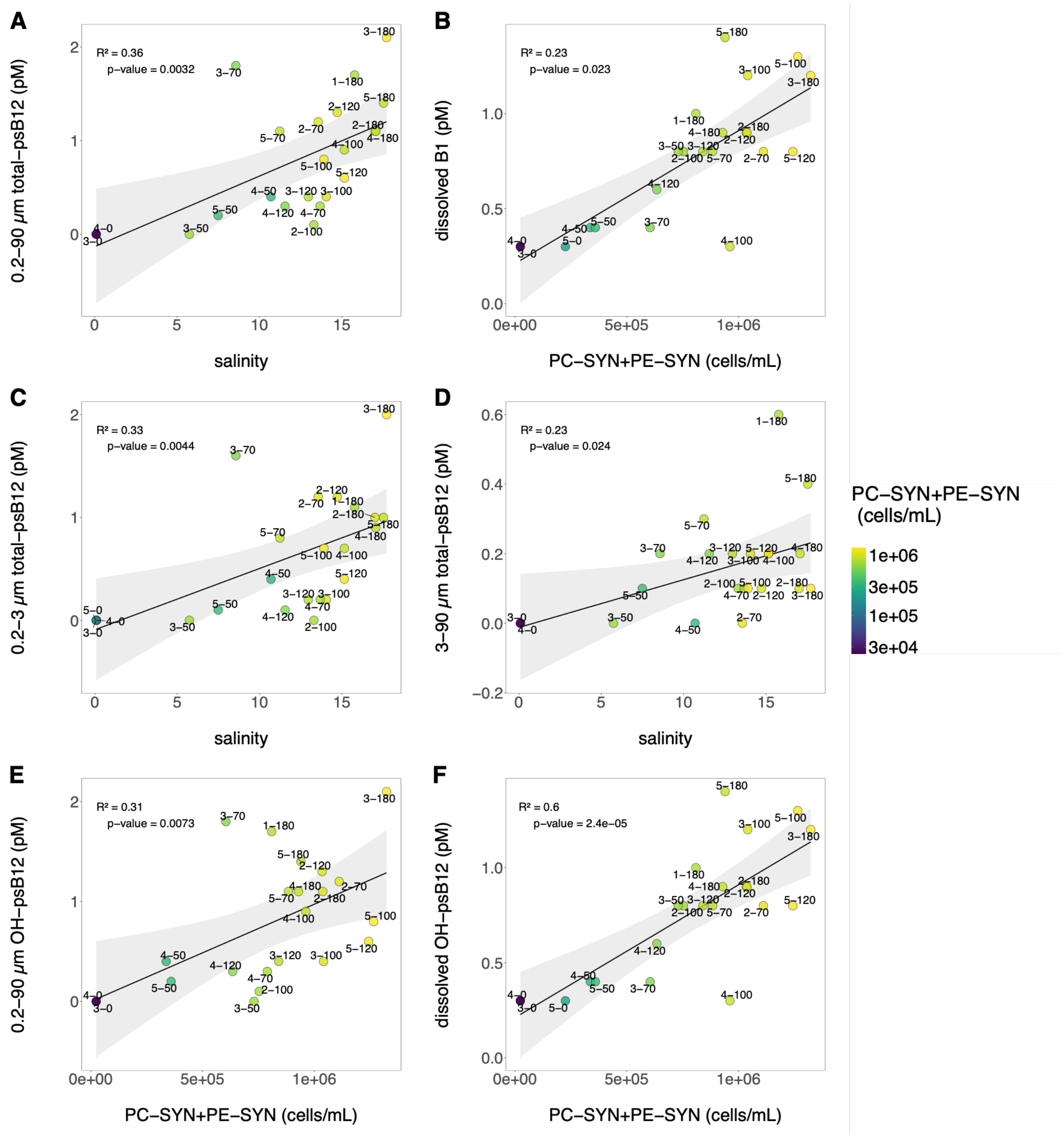


**Fig. S8** Relationship between psB12/B1 concentrations and salinity or picocyanobacterial abundance (PC-SYN+PE-SYN; PC-SYN: *Synechococcus*-like phycocyanin (PC)-rich cells); PE-SYN: *Synechococcus*-like phycoerythrin (PE)-rich cells). Particulate (0.2-90 µm) total psB12 vs. salinity (**A**) and dissolved B1 vs. picocyanobacterial abundance (cells/mL; **B**). Particulate total psB12 from the picoplankton (0.2-3 µm) and the nano-/microplankton (3-90 µm) vs. salinity (**C**, **D**). Particulate (0.2-90 µm) total psB12 and dissolved OH-psB12 vs. picocyanobacterial abundance (cells/mL; **E**, **F**). As peak particulate psB12 concentrations were observed at NRE0 on 25 October, the 3-90 µm measurement was excluded from the regression analysis (**A**, **D**, **E**). Gray shading indicates the 95% confidence interval around the linear regression.

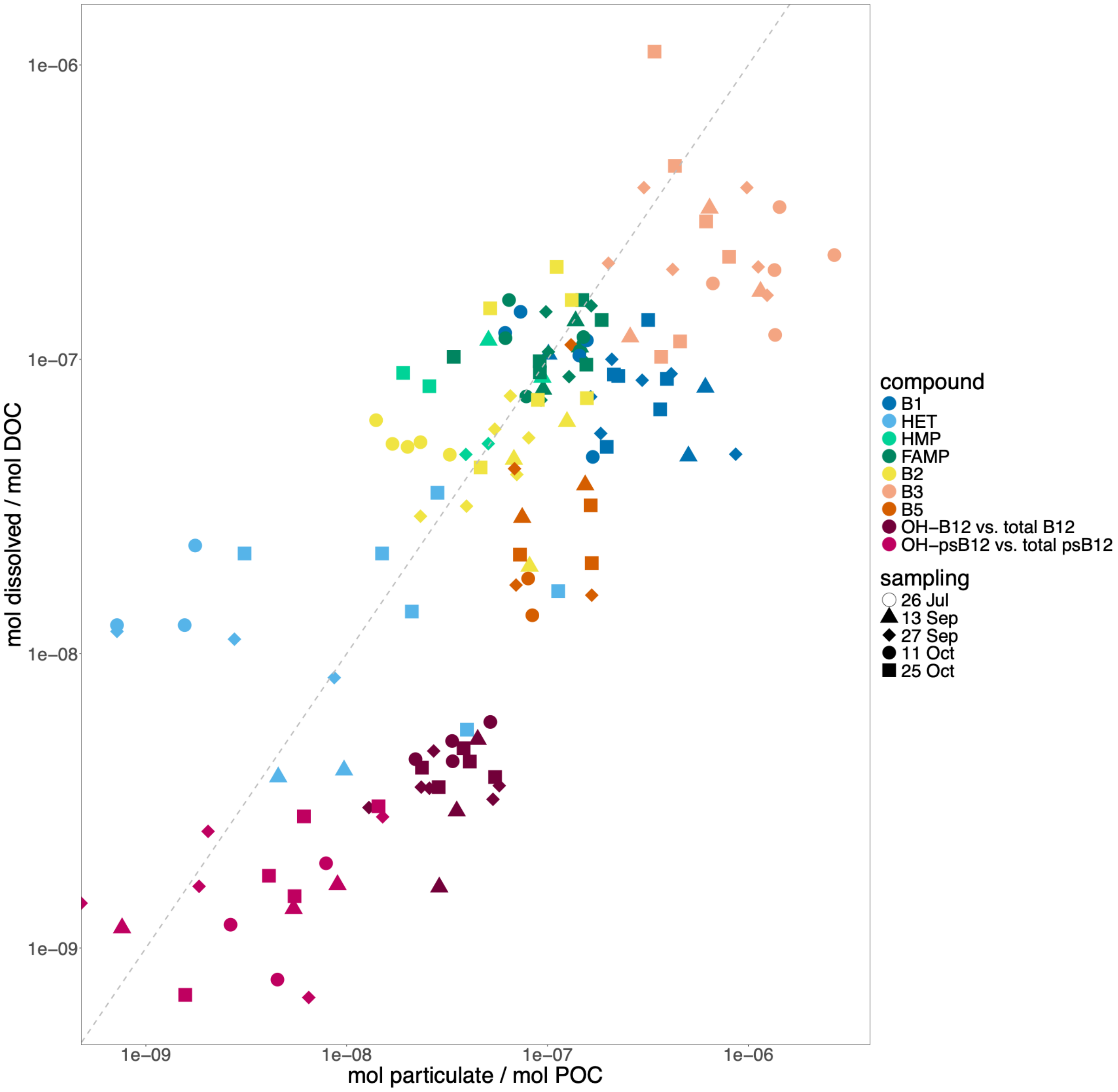


**Fig. S9.** Scatter plot of phase partitioning of B-vitamins and vitamers. Log 10 scale of particulate mole metabolite per mole organic carbon versus dissolved mole metabolite per mole organic carbon. Shape of points indicates sampling time point and color corresponds to metabolite compound. Gray dashed line indicates a 1:1 ratio of particulate to dissolved. Points left of the line indicate an enrichment in the dissolved phase while points right of the line indicate an enrichment in the particulate phase. Only metabolites with measurements in both phases were included, particulate total B12 is the sum of Ado-B12, Me-B12 and OH-B12. Particulate total psB12 is the sum of OH-psB12 and Me-psB12.

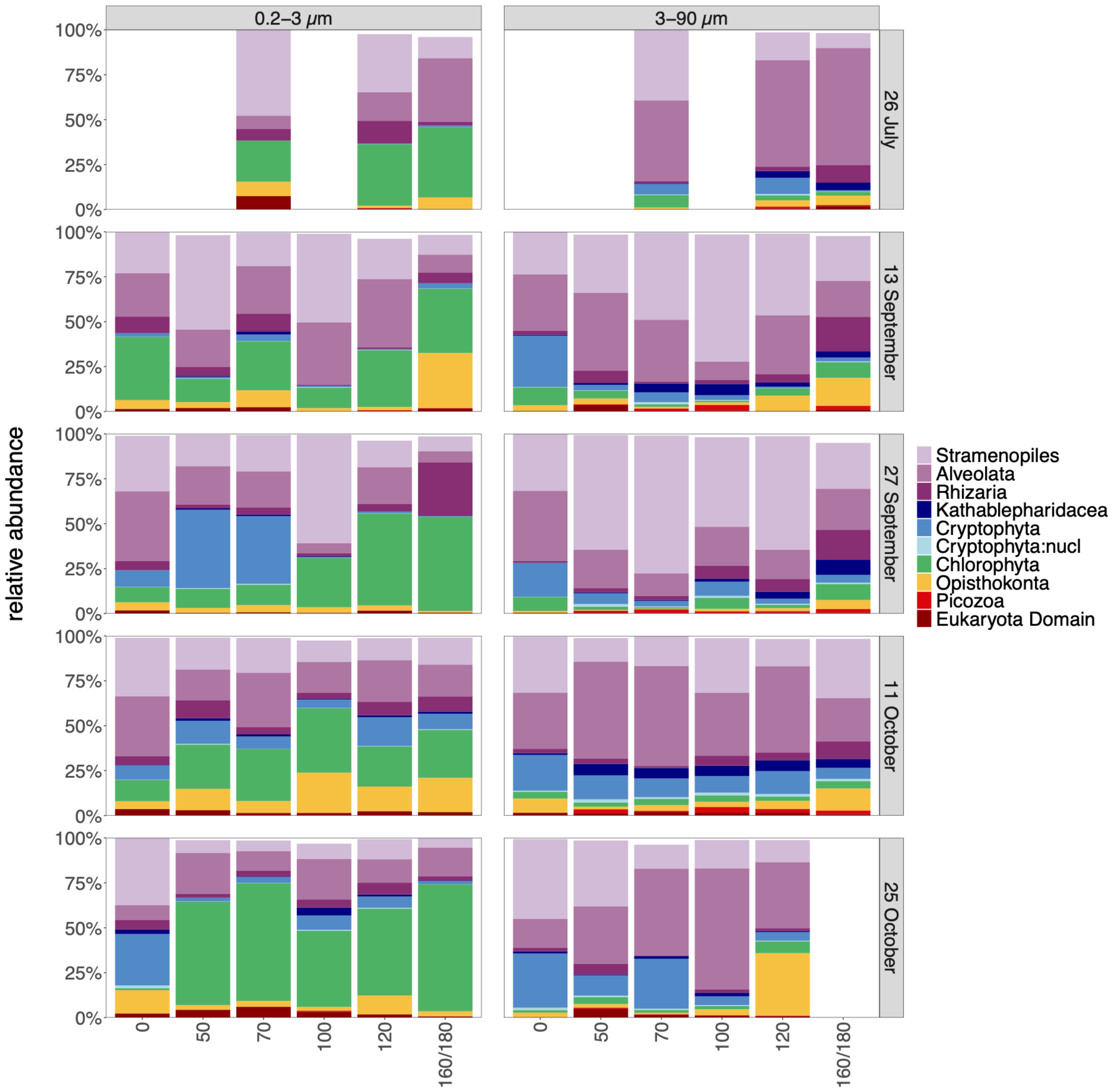


**Fig. S10.** Relative abundance of eukaryotic divisions in picoplankton (0.2-3 µm) and microplankton (3-90 µm) size fraction. Sample NRE180 from the microplankton size fraction was removed due to low sequencing depth.

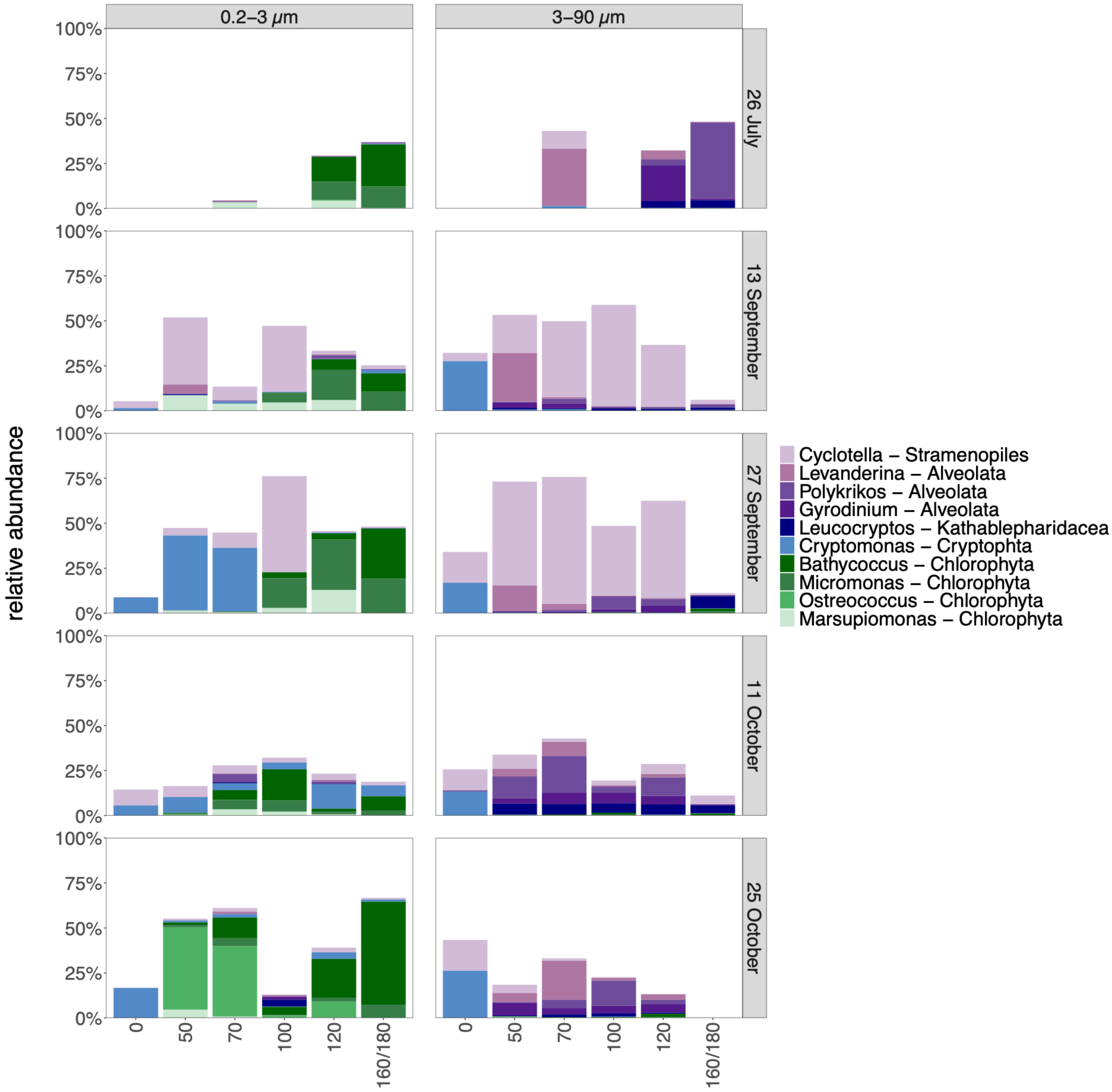


**Fig. S11.** Relative abundance of the top 10 eukaryotic genera that were annotated to a genus level in picoplankton (0.2-3 µm) and microplankton (3-90 µm) size fraction. Sample NRE180 from the microplankton size fraction was removed due to low sequencing depth.

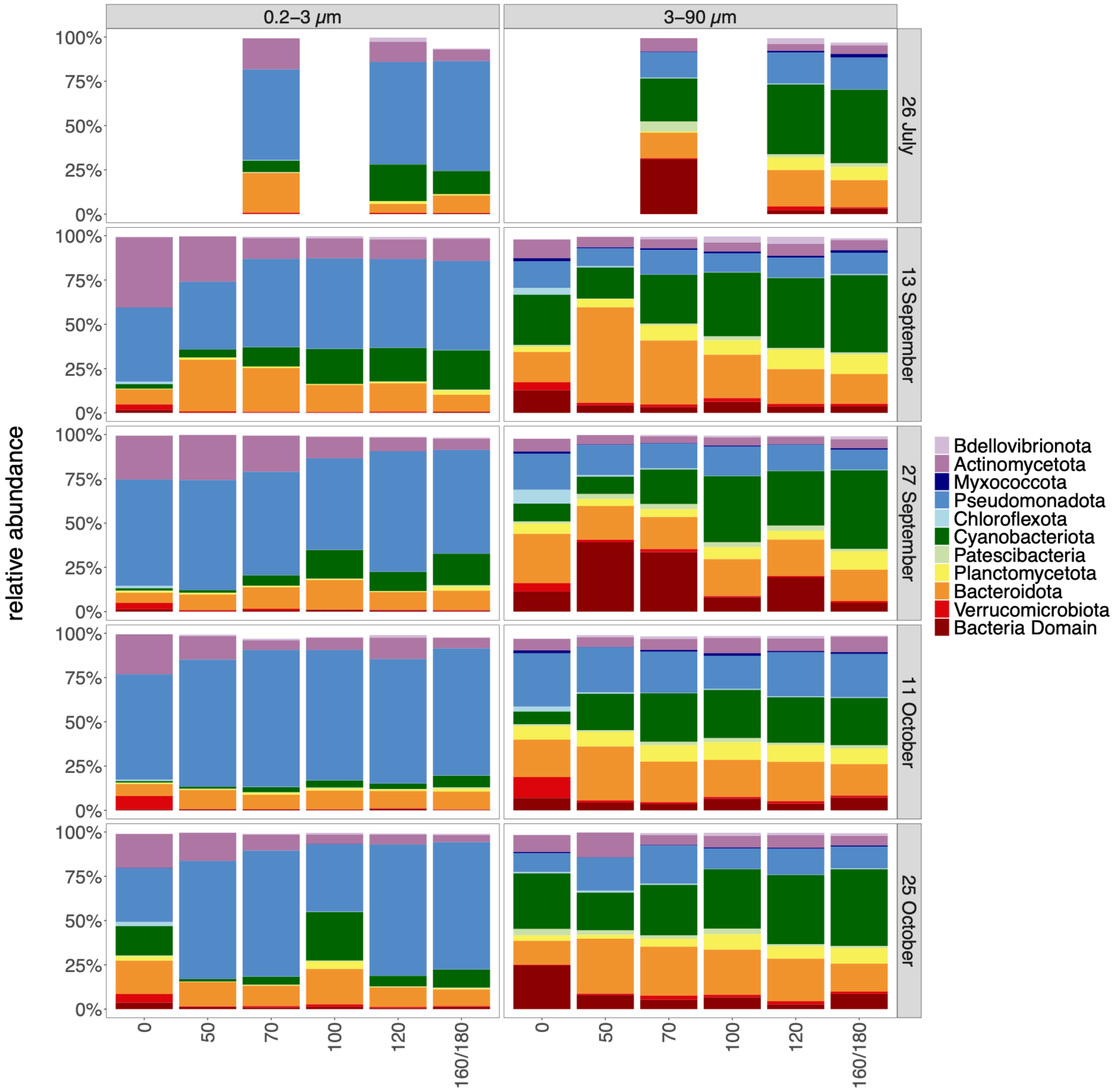


**Fig. S12.** Relative abundance of prokaryotic phyla in picoplankton (0.2-3 µm) and microplankton (3-90 µm) size fraction based on 16S rRNA gen amplicons.

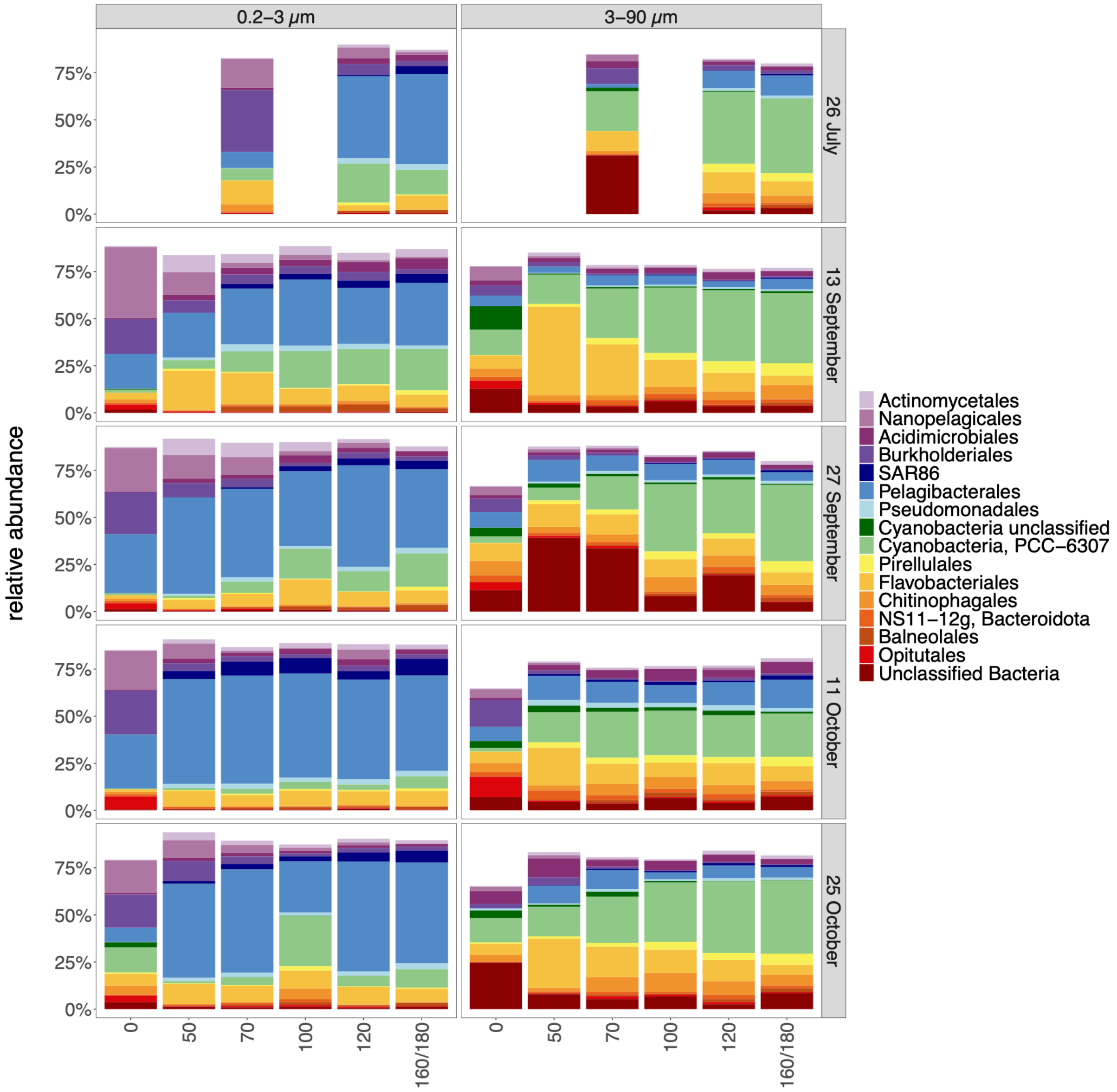


**Fig. S13.** Relative abundance of prokaryotic orders in picoplankton (0.2-3 µm) and microplankton (3-90 µm) size fraction based on 16S rRNA gen amplicons.

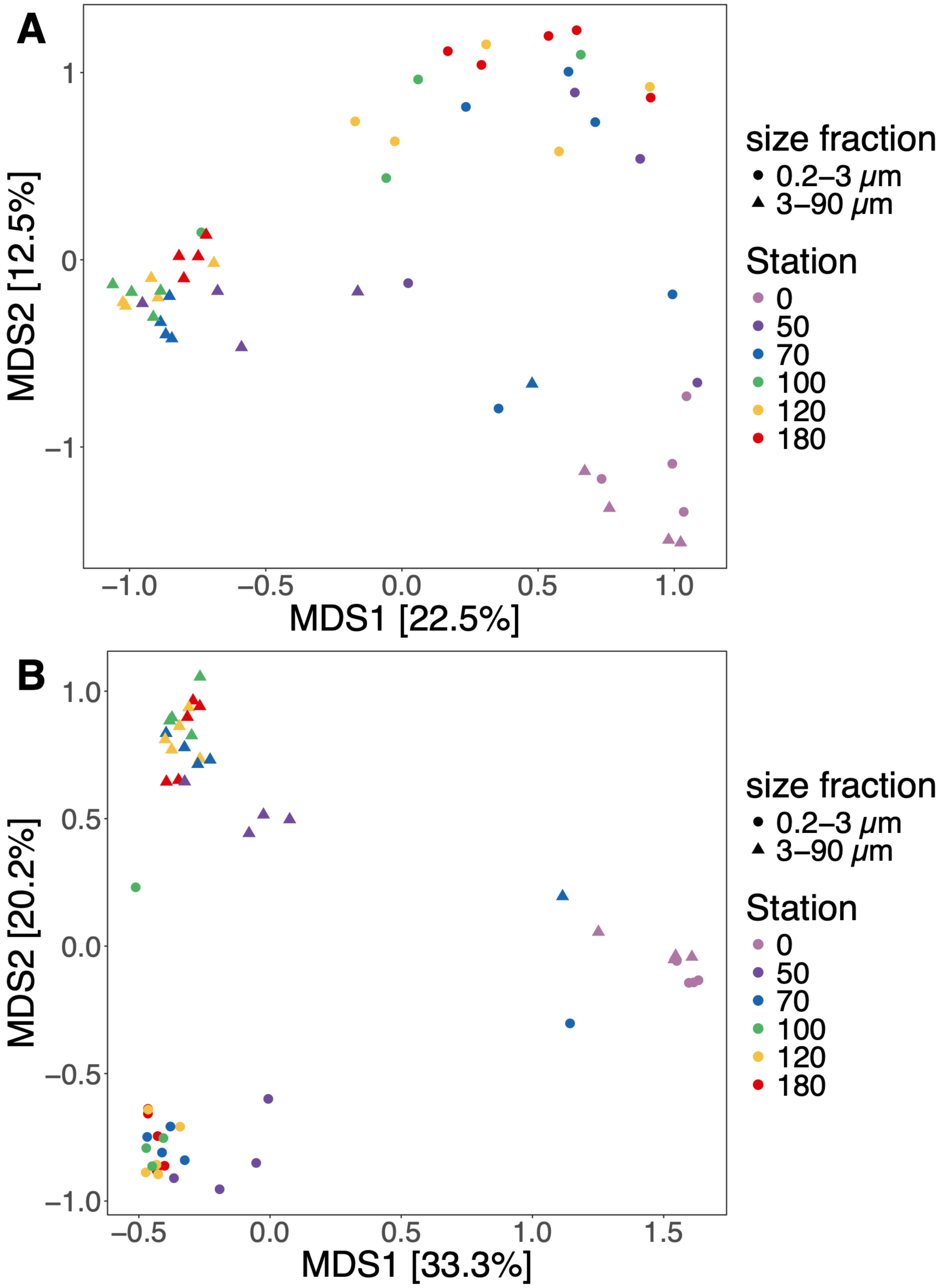


**Fig. S14.** Principal Coordinate Analysis (PCoA) of prokaryotic (**A**) and eukaryotic (**B**) plankton community composition. Abundance tables were Hellinger transformed and Bray-Curtis distance matrixes calculated.

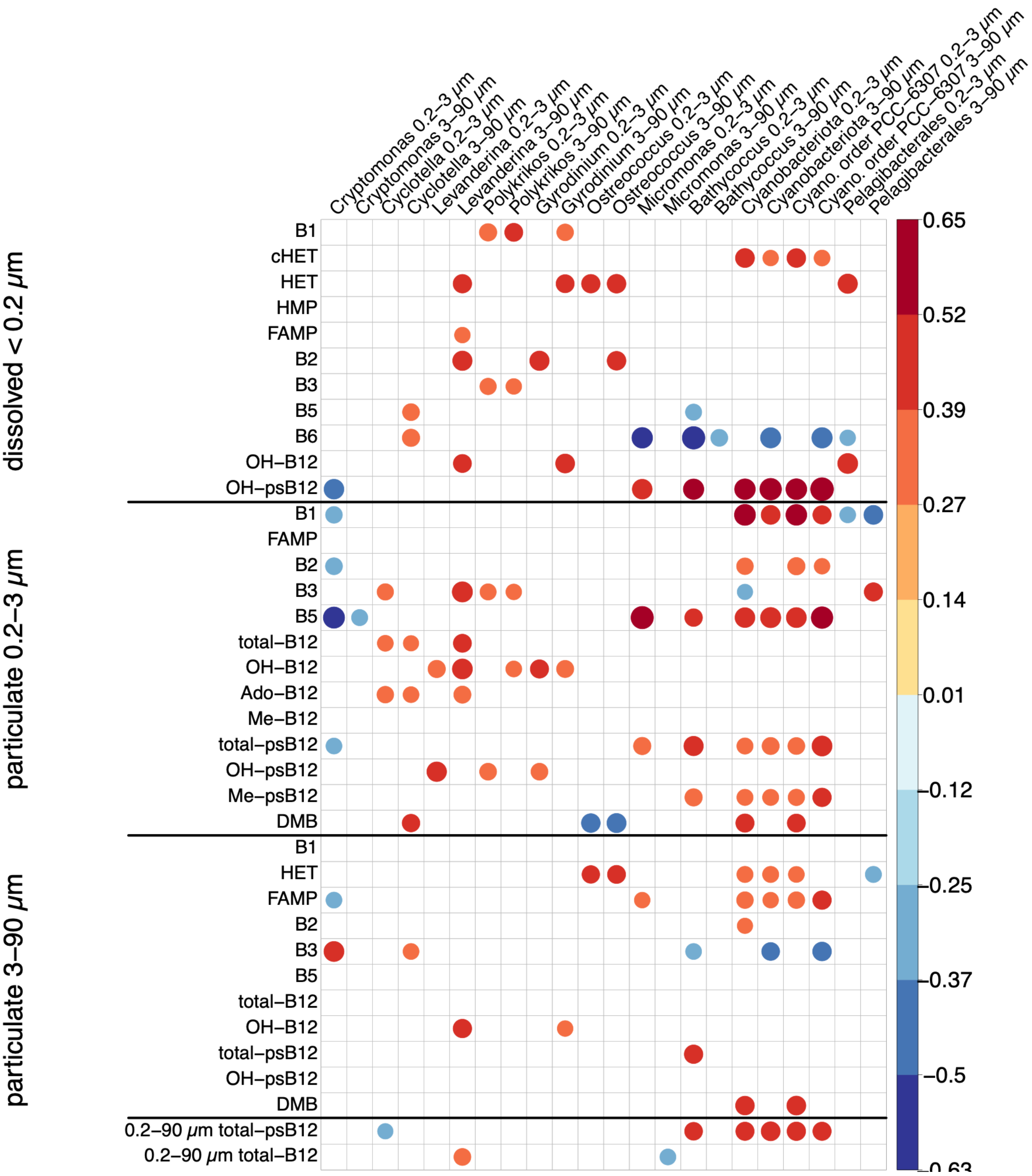


**Fig. S15.** Correlation matrix of significant ($p < 0.05$) Kendall's rank correlations between relative abundance of select eukaryotic and prokaryotic taxa and metabolite concentrations. PCC-6307 is a cyanobacterial (Cyano.) order spanning multiple picocyanobcterial *Synechococcus* -like taxa, including the genera *Vulcanococcus*, *Cyanobium*, *Synechococcus_C*.

### Supporting Tables

**Table S1**. Percent recoveries applied to dissolved values. HET, HMP, FAMP and cHET recoveries from Paerl et al. (2023b). Sample specific B1 recoveries were applied for each sample, mean ± standard deviation shown.

| | **Compound** | low salinity set | high salinity set |
|---|---|---|---|
| Dissolved | B1, thiamin | 32 ±12 | 34 ±13 |
| Dissolved | HET, 4-methyl-5-thiazoleethanol | 89 | 97 |
| Dissolved | HMP, 4-amino-5-hydroxymethyl-2-methylpyrimidine | 100 | 87 |
| Dissolved | cHET, 5-(2-hyrdoxyethyl)-4-methyl-1,3-thiazole-2-carboxilic acid | 8 | 36 |
| Dissolved | FAMP, N-formyl-4-amino-5-aminomethyl-2-methylpyrimidine | 50 | 46 |
| Particulate 0.2 -3.0 µm | B1, thiamin | 45 ±15 | 31 ±27 |
| Particulate 3.0-90 µm | B1, thiamin | 16 ±21 | 33 ±13 |

**Table S2**. PCR conditions and primers for 16S rRNA gene amplification (**A**) and 18S rRNA gene amplification (**B**). 16S rRNA gene primers from Parada et al. (2016) and 18S rRNA gene primers are from Lin et al. (2017) and modified based on TAReuk454FWD1/TAReukREV3 from Stoeck et al. (2010) to avoid mismatch with haptophytes.

**A**

| Step | Temperature (°C) | Duration | Cycles |
|---|---|---|---|
| Initial denaturation | 95 | 3 min | 1 |
| Denaturation | 95 | 45 sec | 25 |
| Annealing | 50 | 45 sec | 25 |
| Extension | 72 | 1 min | 25 |
| Final extension | 72 | 3 min | 1 |

Forward: 5'-GTGYCAGCMGCCGCGGTAA-3'

Reverse: 5'-CCGYCAATTYMTTTRAGTTT-3'

**B**

| Step | Temperature (°C) | Duration | Cycles |
|---|---|---|---|
| Initial denaturation | 95 | 5 min | 1 |
| Denaturation | 95 | 40 sec | 30 |
| Annealing | 58 | 2 min | 30 |
| Extension | 72 | 1 min | 30 |
| Final extension | 72 | 7 min | 1 |

Forward: 5'-CCAGCASCYGCGGTAATTCC-3'

Reverse: 5'-ACTTTCGTTCTTGAT-3'

**Table S3.** Variance explained by RDA models and the individual explanatory variables. Particulate B-vitamin/vitamer measurements have 'p_' as a prefix. DOC: dissolved organic carbon, DO: dissolved oxygen.

| community | variable set | adjR2 | variance | significance | explanatory variables | variance | p-value | significance |
|---|---|---|---|---|---|---|---|---|
| 18S | B-vitamins | 0.306 | 0.471 | *** | p_Niacinamide | 0.058 | 0.001 | *** |
| | | | | | B1 | 0.057 | 0.001 | *** |
| | | | | | p_OH-B12 | 0.045 | 0.001 | *** |
| | | | | | p_FAMP | 0.031 | 0.002 | ** |
| | | | | | OH-psB12 | 0.032 | 0.005 | ** |
| | | | | | B6 | 0.030 | 0.003 | ** |
| | | | | | p_DMB | 0.025 | 0.011 | * |
| | | | | | FAMP | 0.024 | 0.016 | * |
| | | | | | p_B2 | 0.022 | 0.022 | * |
| | | | | | p_B5 | 0.023 | 0.016 | * |
| 18S | environmental | 0.139 | 0.148 | *** | Temperature | 0.038 | 0.006 | ** |
| | | | | | Salinity | 0.069 | 0.001 | *** |
| | | | | | DOC | 0.041 | 0.002 | ** |
| 16S | B-vitamins | 0.420 | 0.247 | *** | B1 | 0.070 | 0.001 | *** |
| | | | | | OH-psB12 | 0.041 | 0.001 | *** |
| | | | | | B6 | 0.021 | 0.011 | * |
| | | | | | p_B5 | 0.018 | 0.026 | * |
| | | | | | p_DMB | 0.046 | 0.001 | *** |
| | | | | | p_Niacinamide | 0.025 | 0.002 | ** |
| | | | | | p_FAMP | 0.035 | 0.001 | *** |
| | | | | | p_OH-B12 | 0.020 | 0.012 | * |
| 16S | environmental | 0.204 | 0.158 | *** | Temperature | 0.026 | 0.014 | * |
| | | | | | Salinity | 0.085 | 0.001 | *** |
| | | | | | DO | 0.025 | 0.014 | * |
| | | | | | Turbidity | 0.022 | 0.044 | * |

**Caption for Supporting Information Data**

**Data S1.** Physical, chemical and biological characteristics of NRE across stations and sampling dates. Abundances of the four small phytoplankton morphotypes were detected with flow cytometry *Synechococcus*-like phycoerythrin (PE)-rich (PE-SYN), *Synechococcus*-like phycocyanin (PC)-rich cells (PC-SYN), picoeukaryotic phytoplankton (PEUK) larger eukaryotic phytoplankton cells (LEUK). https://doi.org/10.11583/DTU.31353040

**Data S2.** Compound specific LC-MS parameters for dissolved (**A**) and particulate samples 0.2-3.0 µm (**B**) and 3-90 µm size-fractions (**C**). Mass over charge (m/z) values for the transitions used for quantification of the compound. The limit of detection (LOD) is calculated as three times the variation of the quality control sample. Limit of quantification (LOQ) is calculated as ten times the variation of the quality control sample. LOD and LOQ are given in fmol. Response of OH-pseudocobalamin and Me-pseudocobalamin assumed to be the same as for OH-cobalamin and Me-cobalamin, respectively. Asterisk (*) indicates values between LOD and LOQ were verified by the batch per batch method, otherwise only values above LOQ were quantified. https://doi.org/10.11583/DTU.31353040

**Data S3**. Concentrations of B-vitamins and vitamers present in the dissolved (**A**), small particulate (0.2-3 µm) size-fraction (**B**), and large (3-90 µm) particulate size-fraction (**C**) in the near-surface waters of the Neuse River Estuary across time points and stations sampled. Number of biological replicates for each sample for which data is available is provided in the column labelled n, as some replicates were lost during sample processing or measurement (see Methods). Not all metabolites were detected in all biological replicates as, therefore the number of biological replicates, on which the measurement is based, is provided for each metabolite. If the metabolite was detected in minimum three replicates the standard deviation (sd) is listed. LOD: measurements were below limit of detection. LOQ: measurements were below limit of quantification. Compounds specific LODs and LOQs are provided per dataset and matrix grouping in Supporting Information **Data S2**.
https://doi.org/10.11583/DTU.31353040
